# Phagocytosis via complement receptor 3 enables microbes to evade killing by neutrophils

**DOI:** 10.1101/2022.09.01.506228

**Authors:** Asya Smirnov, Kylene P. Daily, Mary C. Gray, Stephanie A. Ragland, Lacie M. Werner, M. Brittany Johnson, Joshua C. Eby, Erik L. Hewlett, Ronald P. Taylor, Alison K. Criss

**Affiliations:** Department of Microbiology, Immunology, and Cancer Biology, University of Virginia School of Medicine; Division of Infectious Diseases and International Health, Department of Medicine, University of Virginia School of Medicine; Department of Biochemistry and Molecular Genetics, University of Virginia School of Medicine

**Keywords:** *Neisseria gonorrhoeae*, neutrophil, phagocytosis, integrin, phagocytosis, CR3, complement, *Mycobacterium smegmatis*

## Abstract

Complement receptor 3 (CR3; CD11b/CD18; α_m_β_2_ integrin) is a conserved phagocytic receptor. The active conformation of CR3 binds the iC3b fragment of complement C3 as well as many host and microbial ligands, leading to actin-dependent phagocytosis. There are conflicting reports about how CR3 engagement affects the fate of phagocytosed substrates. Using imaging flow cytometry, we confirmed that binding and internalization of iC3b-opsonized polystyrene beads by primary human neutrophils was CR3-dependent. iC3b-opsonized beads did not stimulate neutrophil reactive oxygen species (ROS), and most beads were found in primary granule-negative phagosomes. Similarly, *Neisseria gonorrhoeae* (Ngo) that does not express phase-variable Opa proteins suppresses neutrophil ROS and delays phagolysosome formation. Here, binding and internalization of Opa-deleted (Δopa) Ngo by adherent human neutrophils was inhibited using blocking antibodies against CR3 and by adding neutrophil inhibitory factor, which targets the CD11b I-domain. Neutrophils did not produce detectable amounts of C3 to opsonize Ngo. Conversely, overexpressing CD11b in HL-60 promyelocytes enhanced Δopa Ngo phagocytosis, which required CD11b I domain. Phagocytosis of Ngo was also inhibited in mouse neutrophils that were CD11b-deficient or treated with anti-CD11b. Phorbol ester treatment upregulated surface CR3 on neutrophils in suspension, enabling CR3-dependent phagocytosis of Δopa Ngo. Neutrophils exposed to Δopa Ngo had limited phosphorylation of Erk1/2, p38, and JNK. Neutrophil phagocytosis of unopsonized *Mycobacterium smegmatis*, which also resides in immature phagosomes, was CR3-dependent and did not elicit ROS. We suggest that CR3-mediated phagocytosis is a silent mode of entry into neutrophils, which is appropriated by diverse pathogens to subvert phagocytic killing.

## INTRODUCTION

As first responders to infection and sterile inflammation, neutrophils have a dynamic capacity for phagocytosis of both foreign and host material (1). How this material is recognized by host receptors influences the extent of phagocytosis and the subsequent fate of the material. Phagocytosis of substrates can proceed through opsonization by soluble components such as antibody and complement, and can also be independent of opsonization by interaction with neutrophil receptors that directly recognize the material. After phagocytosis, the nascent phagosome fuses with cytoplasmic granules to form a mature phagosolysosome, and assembly of NADPH oxidase on the phagosome generates antimicrobial reactive oxygen species (ROS), a hallmark of neutrophil activation (2, 3). Fusion of secondary/specific granules to phagosomes or to the plasma membrane releases pro-forms of cathelicidin antimicrobial peptides, the nutritional immunity protein lactoferrin, and the cytochrome_b558_ component of NADPH oxidase, while primary/azurophilic granule fusion releases serine proteases such as neutrophil elastase, α-defensins, and myeloperoxidase (2). The concerted action of these factors in the phagolysosome mediates killing of microbes and digestion of the phagocytosed material (1). Granule mobilization, ROS production, and kinase cascades that lead to production and release of proinflammatory cytokines are modulated by the signals emanating from phagocytic receptors, as well as through co-receptors that recognize the material but are not themselves phagocytic (4).

Neutrophils are characterized by the abundant surface expression of complement receptor 3 (CR3; α_m_β_2_; CD11b/CD18). CR3 is an integrin heterodimer that is critical for the chemotaxis and phagocytic activity of neutrophils and macrophages (5). On naïve phagocytes CR3 is in an inactive conformation. Complex inside-out and outside-in signals place CR3 in an active, ligand binding-proficient state, with the CD18 cytoplasmic tail interacting with the F-actin cytoskeleton via talin, vinculin, and kindlin (reviewed in (6)). Although the canonical ligand for CR3 is the complement fragment iC3b, CR3 binds a diverse array of ligands that are both host-and microbial-derived, including extracellular matrix proteins, the cathelicidin LL-37, fungal β-glucan, and bacterial toxins, lipopolysaccharide, and adhesins (7–15). Binding to iC3b and many other ligands is predominantly mediated by the I-domain, an extension that is found in leukocyte CD11 proteins relative to other α integrins (5). Overlapping with, but distinct from the CD11b I-domain is the metal ion-dependent adhesion site (MIDAS) that also promotes ligand binding (5). While the ability of CR3 to facilitate phagocytosis is well accepted, there are differing reports concerning whether CR3 engagement leads to phagolysosome formation and ROS production by neutrophils (reviewed in (16)).

Given the central role of neutrophils in recognition of and response to microbial challenge, it is not surprising that some pathogens make use of mechanisms to resist phagocytic killing by neutrophils. These mechanisms include evasion of opsonophagocytosis by preventing antibody or complement deposition, expression of antiphagocytic surface components like capsules, use of toxins or bacterially-injected proteins to block phagosome-granule fusion, and resistance to neutrophil antimicrobial components (reviewed in (17)). One pathogen that evades neutrophil-mediated clearance is the bacterium *Neisseria gonorrhoeae* (Ngo), which causes the sexually transmitted disease gonorrhea (18). In symptomatic gonorrhea, neutrophils are recruited in abundance to mucosal sites of infection.

However, viable, infectious Ngo are recovered from the resulting neutrophil-rich purulent exudates, indicating Ngo has strategies to resist clearance by neutrophils (19).

The major determinant of the fate of Ngo inside neutrophils is how it is phagocytosed, which is influenced by the physiological state of the neutrophil (20). The predominant driver of nonopsonic interaction of Ngo with neutrophils is the family of opacity-associated (Opa) surface-exposed proteins (21–23). Each of the 10-14 *opa* genes in Ngo is independently phase-variable, conferring an extensive capacity for surface variation in the bacterial population. Most Opa proteins bind human carcinoembryonic antigen-related cell adhesion molecules (CEACAMs), of which CEACAMs 1, 3, and 6 are expressed by neutrophils (24). Opa-expressing (“Opa+”) Ngo that bind the granulocyte-restricted, immunotyrosine activation motif (ITAM)-bearing CEACAM3 is rapidly phagocytosed into mature phagolysosomes and stimulates reactive oxygen species (ROS) production (25–27). In contrast, unopsonized Opa-negative Ngo is not phagocytosed by neutrophils in suspension and does not stimulate ROS production (20, 28, 29). We have reported that adherent, interleukin-8 treated human neutrophils can phagocytose Opa-negative Ngo, although they do so more slowly than when they internalize Opa+ bacteria. Moreover, their phagosomes exhibit delayed fusion with primary granules, enabling intracellular survival (30–32). IgG-opsonized Ngo phenocopies CEACAM3-binding bacteria, while serum-opsonized bacteria do not stimulate ROS production or phagolysosome formation (31). The mechanism by which adherent but not suspension neutrophils phagocytose unopsonized, Opa-negative Ngo has not been identified.

In this study we define CR3-mediated phagocytosis as a mechanism of “silent entry” into neutrophils, which does not trigger ROS production, early phagolysosome formation, or intracellular proinflammatory signaling cascades. These outcomes are shown for three unrelated substrates: iC3b-opsonized beads, *Mycobacterium smegmatis*, and Opa-negative Ngo, which we found exploits CR3 for phagocytosis by adherent neutrophils. Suspension neutrophils gained the capacity for phagocytosis of Opa-negative Ngo when CR3 was ectopically activated, and overexpression of CD11b enabled phagocytosis of Opa-negative Ngo by HL-60 human promelocytes. From these results, we posit that neutrophils use CR3 for silent phagocytosis of material with limited cellular activation. This process is exploited by diverse pathogens to survive confrontation with these otherwise antimicrobial cells.

## MATERIALS AND METHODS

### Chemicals, reagents, and antibodies

All chemicals and reagents were from ThermoFisher Scientific unless otherwise noted. Antibody sources and reactivities are provided in **Table 1**.

**Table 1.**
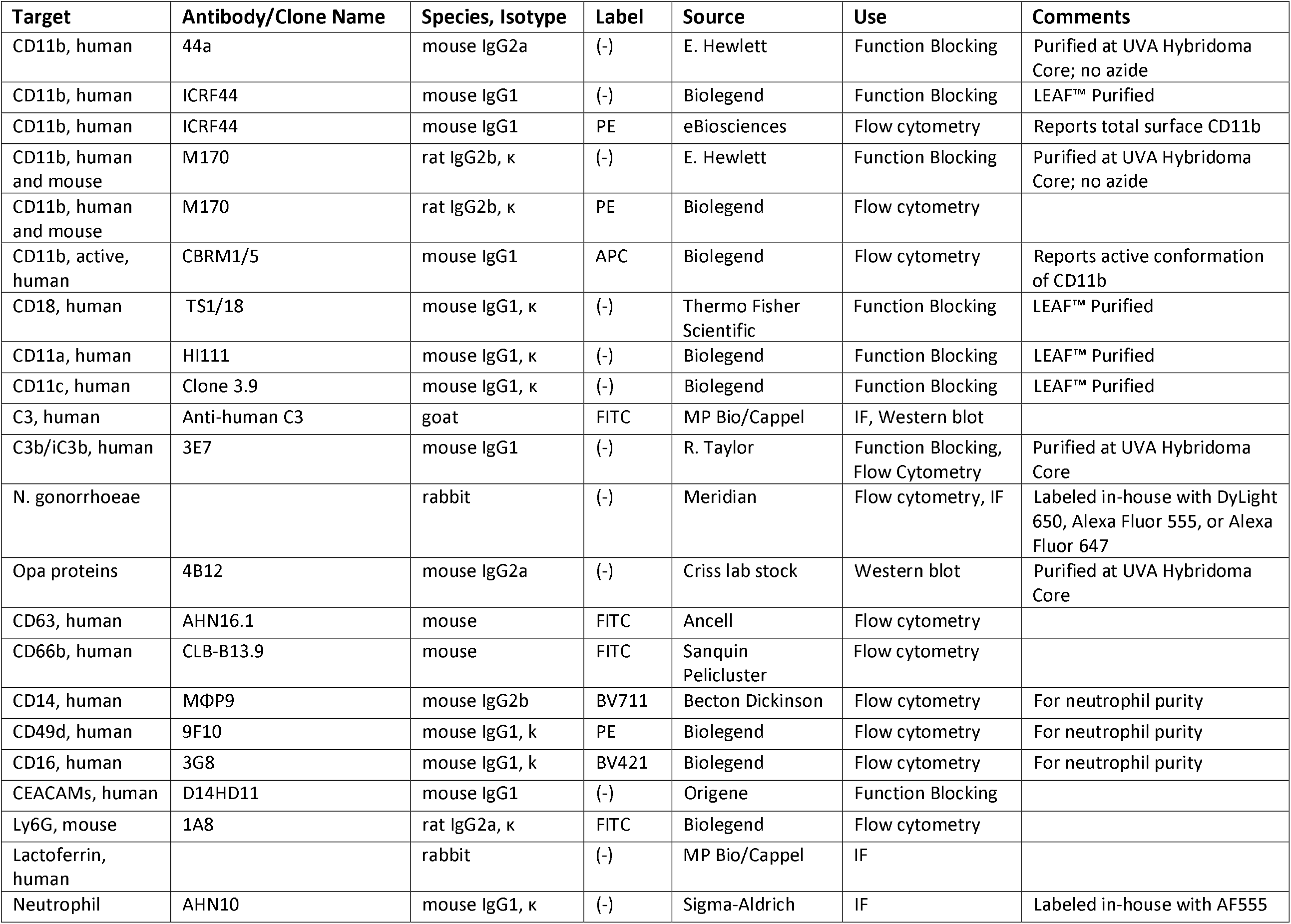
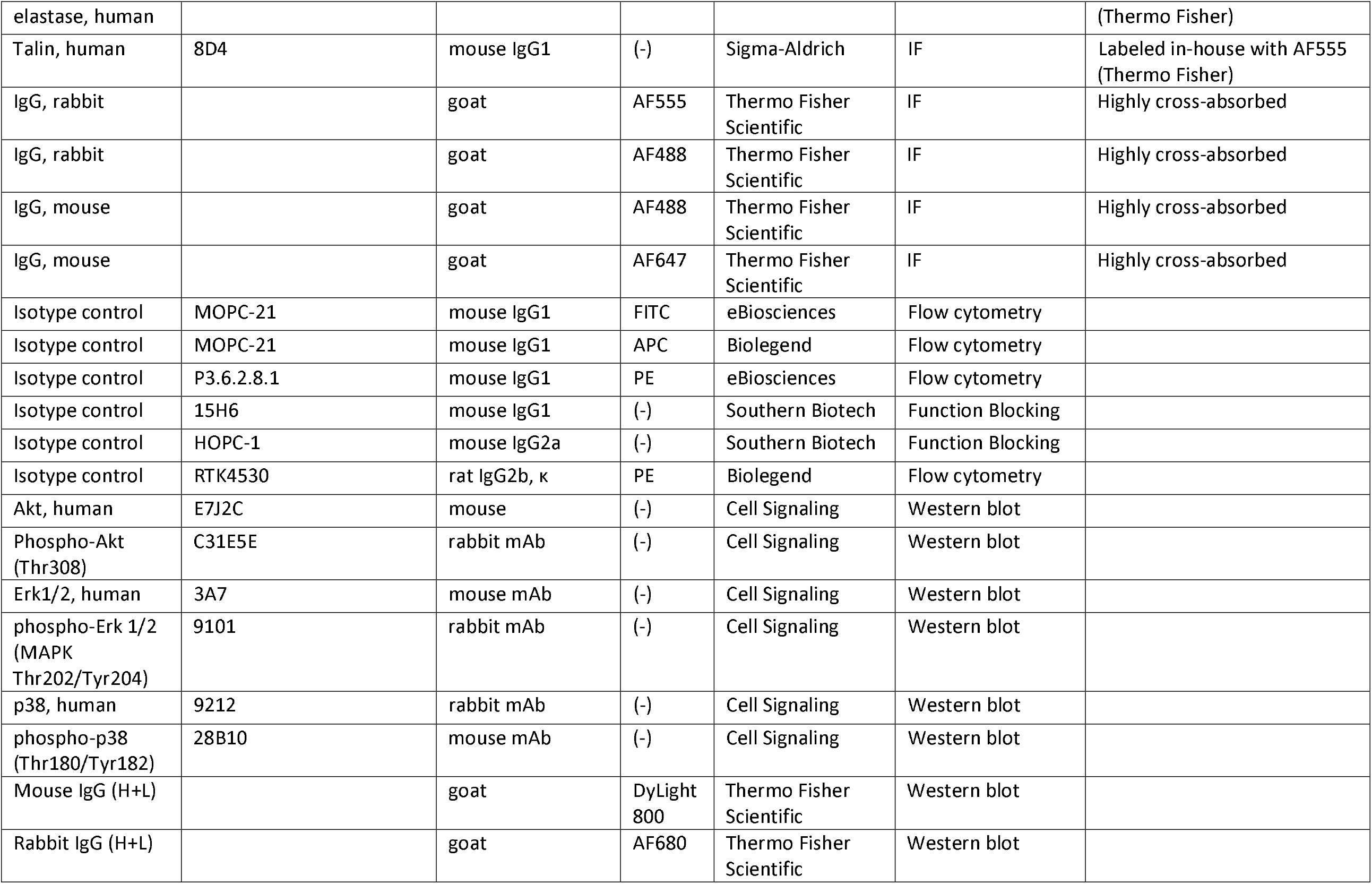
Antibodies used in this study.

### Bacterial strains and growth conditions

This study used the constitutively piliated Δopa (Opaless; *ΔopaA-K*) and isogenic constitutively OpaD-expressing (OpaD+) Ngo of strain background FA1090 (33)). Other strain backgrounds used were piliated, *recA6* FA1090 RM11.2 (34), constitutively piliated derivative of MS11 VD300 (35), and piliated 1291 (gift of M. Apicella, Univ. of Iowa). Phenotypically translucent colonies from these three Opa-variable strain backgrounds were used for experiments. The absence of Opa expression in these derivatives was confirmed by immunoblotting with the 4B12 anti-Opa antibody (36).

Ngo was grown on solid GCB media containing Kellogg’s supplement I and II (37) for 16-18 hr at 37°C, 5% CO_2_. Viable, exponential-phase predominantly piliated (except the Δ*pilE* and Δ*pilQ* mutants) bacteria were grown in rich liquid medium with three sequential dilutions (38). For opsonization, Ngo was incubated with gentle rotation in 10% pooled normal human serum (Sigma) for 20 min at 37°C as described previously (32). Complement deposition on the Ngo surface was visualized by immunofluorescence using a fluorescein-conjugated antibody against human C3.

*M. smegmatis* (Msm) strain mc2 155 was from G. Ramakrishnan (Univ. of Virginia), and an mCherry-expressing derivative (mgm1732; generated by transformation of mc2 155 with the pCherry3 mCherry expression plasmid (39, 40)) was from C. Stallings (Washington Univ.). Bacteria were maintained on LB agar with 50% glycerol and 40% dextrose (with 50 µg ml^−1^ hygromycin for mgm1732). For experiments, bacteria were inoculated in LB liquid medium with 50% glycerol, 40% dextrose, and 0.25% Tween 80 (Sigma) (with 50 µg ml^−1^ hygromycin for mgm1732) and shaken at 150 rpm at 37 °C for 48 hr.

Where indicated, bacteria were labeled with 5 µg ml^−1^ carboxylfluorescein diacetate succinimidyl ester (CFSE) or Tag-IT Violet™ proliferation and cell tracking dye (Biolegend) in PBS containing 5 mM MgSO_4,_ at 37°C for 20 min in the dark, then pelleted and washed before use in experiments.

### Ngo mutant construction

*ΔpilE:* Overlap extension PCR was used to replace most of the *pilE* open reading frame with a PvuII restriction site. Using genomic DNA from FA1090 that is nonvariable for the 1-81-S2 *pilE* sequence and has deletions in *opaB, opaE, opaG*, and *opaK* (33), the 5’ end of *pilE* and flank was amplified with primers PilE-RA 5’-TTGGGCAACCGTTTTATCCG-3’ and PilE-FA 5’-GCCTAATTTGCCTTAGGTGGCAGA<u>CAGCTG</u>TTATTGAAGGGTATTCATAAAATTACTCC-3’, the 3’ end of *pile* and flank was amplified with primers PilE-RB 5’-GGCGTCGTCTTTGGCGA-3’ and PilE-FB 5’-GGAGTAATTTTATGAATACCCTTCAATAA<u>CAGCTG</u>TCTGCCACCTAAGGCAAATTAGGC-3’ (PvuII site underlined), and the two products were combined for overlap extension PCR using PilE-FA and PilE-RB. The resulting 1.5 kb product was spot transformed into Δopa Ngo (41), and P-colonies were phenotypically selected and streak purified. The presence of the deletion was confirmed by PCR using primers PILRBS and PilE-RA, and Southern blot of XhoI and ClaI-digested genomic DNA with a digoxygenin-labeled *pilE* probe, amplified from the piliated parent.

*pilT:* The *pilT* gene and flanking sequence from the FA1090 chromosome was amplified by PCR with primers PilT-F 5’-CATTGAGGTCGGCAAGCAGC-3’ and PilT-R 5’-GCATCTTTACCCAGCGCGAAAT-3’ and cloned into pSMART HC Kan (Lucigen). The cloning mix was transformed into TOP10 *E. coli*, transformants were selected on 50 µg ml^−1^ kanamycin, and the presence of the insert was confirmed by PCR and sequencing (Genewiz). The plasmid was purified from *E. coli* by miniprep (Qiagen), and the internal 504 bp of *pilT* was removed by digestion with AccI (New England Biolabs). The plasmid was religated with T4 DNA ligase (New England Biolabs) and transformed into TOP10 *E. coli*. The presence of the deletion was confirmed by PCR and sequencing. The Δ*pilT* plasmid was spot transformed into Δopa Ngo, and transformants were selected based on the characteristic jagged edge colony morphology of *pilT* mutant Ngo. Transformants were passaged on GCB, and the presence of the deletion was determined by product size following PCR amplification with ΔPilT-F 5’-TTACCGACTTACTCGCCTTCG-3’ and ΔPilT-R 5’-CGATTGCAGCGATTGGTCC-3’ and confirmed by DNA sequencing.

*pilQ:* Δopa Ngo was spot transformed with a plasmid carrying the *pilQ* gene interrupted by a chloramphenicol resistance cassette (from H. Seifert, Northwestern Univ.) (42). Transformants were selected on 0.5 µg ml^−1^ chloramphenicol and passaged sequentially three times on chloramphenicol-GCB. The presence of *pilQ::cat* was confirmed by P-colony morphology, PCR using ΔpilQ-F 5’-CGCAACCGCCGCCTTTCA-3’ and ΔpilQ-R 5’-CAGCCTGCGCTATTGATGC-3’, and Southern blot using a *pilQ* digoxygenin-labeled probe.

*pglA, pglD:* Genomic DNA from *pglA::kan* and *pglD::kan* in strain background 1291 (from J. Edwards, Nationwide Children’s) was transformed into Δopa Ngo, followed by two more rounds of backcrossing from confirmed transformants. In all cases, transformants were selected on GCB containing 50µg mL^−1^ kanamycin, and the presence of the mutated gene confirmed by PCR and sequencing (*pglA:* ΔpglA-F 5’-ACAACAGTCGCATCCAGCAT-3’ and ΔpglA-R 5’-CGGGGATTCCAAGGTTCGAT-3’; *pglD:* ΔpglD-F 5’-ATCTGGAAACCCTGATCGCC’3’ and ΔpglD-R 5’-TTCGCATACCCAAAGCAGGT-3’).

### Human neutrophil purification

Peripheral venous blood was drawn from healthy human subjects with informed consent, in accordance with a protocol approved by the University of Virginia Institutional Review Board for Health Sciences Research (#13909). Neutrophils were purified by dextran sedimentation followed by Ficoll separation and hypotonic lysis of residual erythrocytes as in (38).

Neutrophil content in suspension was >90% as monitored by phase-contrast microscopy and confirmed by flow cytometry (CD11b+/CD14-/ CD49d^low^/CD16^high^). All replicate experiments were conducted using neutrophils from different subjects.

### Bacterial association with and internalization by adherent primary human neutrophils

Neutrophils were suspended in RPMI (Cytiva) + 10% heat-inactivated fetal bovine serum (HyClone) (hereafter referred to as “infection medium”) containing 10 nM human IL-8 (R&D Systems) and left to adhere to tissue culture-treated plastic coverslips (Sarstedt) for 30 min. Neutrophils were treated with blocking antibodies or matched isotype control (20 µg ml^−1^; see **Table 1**), or recombinant *A. caninum* NIF protein (R&D Systems; 0.5 µg ml^−1^) for 20 min. Fluorescently labeled Ngo was then added to neutrophils by centrifugation at 400 xg for 4 min at 12 °C. The supernatant was discarded to remove unbound bacteria, pre-warmed infection medium was added, and the cells and bacteria were coincubated for 1 hr at 37 °C and 5% CO_2_ (38). Cells were fixed with 2% paraformaldehyde (PFA) (Electron Microscopy Sciences) for 10 min on ice, collected into microfuge tubes using a cell scraper, and washed (600 x g, 4 min DPBSG washes x 3). Non-specific antibody binding was blocked with 10% normal goat serum for 10 min followed by staining with anti-Ngo antibody coupled to DyLight™ 650 (ThermoFisher).

Cells were analyzed by imaging flow cytometry using ImageStream^X^ Mark II operated by INSPIRE software (Luminex). Bacterial association with neutrophils was measured using IDEAS 6.2 Software (Luminex) as previously described (43). Briefly, the percent of neutrophils with associated Ngo was calculated as the percent of CFSE+ or Tag-IT Violet+ cells, and the percent of neutrophils with internalized Ngo was calculated using spot count and a step gate function as the percent of cells with ≥ 1 CFSE+ or Tag-IT Violet+ only bacterium. For *M. smegmatis-*infected neutrophils, extracellular bacteria were not identified by differential fluorescence. Instead, intracellular bacteria were defined as those whose entirety was within a mask that was eroded by 4 pixels from the neutrophil periphery, as marked by staining for surface CD11b. The single cell population was identified as the population of cells with medium area and medium to high aspect ratio. Cell morphology was visually verified within the gate boundaries.

### Phagocytosis of opsonized sheep red blood cells by adherent primary human neutrophils

Sheep red blood cells (sRBCs) (MP Biomedicals) were opsonized with iC3b in gelatin veronal buffer, using anti-sRBC IgM and C5-deficient human serum as in (44). Complement deposition on sRBCs was confirmed by using fluorescein-conjugated rabbit anti-human C3 antibody (10 µg mL^−1^, 30 min incubation at 37 °C).

Adherent, IL-8 treated primary human neutrophils, incubated in infection medium, were exposed to 150 ng mL^−1^ phorbol myristate acetate (PMA; Sigma) for 5 min at 37 °C to activate CR3. Neutrophils were incubated with blocking antibodies or isotype control (20 mg ml^−1^, 20 min), then exposed to 10^6^ opsonized sRBCs in infection medium at 37 °C for 10 min. Plates were centrifuged at 600 x g for 4 min at 12°C and then put on ice to stop phagocytosis. Media was removed, extracellular sRBCs were lysed with endotoxin free water (ICUMedical) for 1 min, and neutrophils were incubated in ice cold methanol for 10 min. Cells were washed once with water and mounted on slides for imaging. The phagocytic index is reported as the number of intracellular opsonized sRBCs per 100 neutrophils.

### Ngo infection of human neutrophils in suspension

Primary human neutrophils (3×10^6^) were suspended in Dulbecco’s PBS without calcium or magnesium (Gibco) with 0.1% glucose (DPBSG-). Neutrophils were left untreated or exposed to 200 nM PMA for 10 min. After treatment with blocking antibodies or isotype control (20 mg ml^−1^), unlabeled or CFSE-labeled Ngo was added to the neutrophils at multiplicity of infection = 1 unless otherwise indicated and incubated at 37 °C for 1 hr with slow rotation. Cells were stained with Zombie Violet viability dye (BioLegend), fixed with 1% paraformaldehyde (PFA) for 10 min, and analyzed by imaging flow cytometry as above. Surface presentation of total (ICRF44) and active (CBRM1/5) CD11b was analyzed by flow cytometry.

### Ngo infection of HL-60 cells

HL-60 cells expressing either empty vector, human CD11b full length, or CD11b lacking I-domain (from V. Torres, NYU; (8)) were incubated with CFSE+ Ngo at the indicated MOI at 37°C for 1 hr, followed by anti-Ngo antibody coupled to DyLight™ 650. The percent of CFSE+ HL-60 cells with intracellular Ngo (≥ 1 CFSE only spot) was measured by imaging flow cytometry using a step gate as for primary human neutrophils.

### *N. gonorrhoeae* association with and internalization by mouse neutrophils

Experiments with mice were approved by the Animal Care and Use Committee at the University of Virginia and supported exclusively by internal funds at the University of Virginia. The tibiae, femurs, and vertebrae were harvested from 4-8 week old, male WT (#000664) and *cd11b*^−/-^ (#003991, B6.129S4-*Itgam*^tm1myd^/J) C57BL/6J mice (Jackson Laboratories). Male mice were used as they were available at the time these experiments were initiated; there are no reports to suggest that CR3 on neutrophils from male and female mice would respond differently. Bone marrow was extracted by mortar and pestle, and neutrophils wee purified using an antibody-based negative selection kit (Stem Cell Technologies). Purified neutrophils were >84% viable (using Zombie Violet, Biolegend) and >80% Ly6G^hi^CD11b^hi^. Neutrophils purified from CD11b^−/-^ mice were confirmed to be Ly6G^hi^CD11b^neg^ by flow cytometry.

Plastic tissue culture coverslips were coated with normal mouse serum (50% diluted in DPBS with CaCl_2_ and MgCl_2_) (Sigma) for 1 hr at 37°C with 5% CO_2_, and residual serum was removed by aspiration. Mouse neutrophils were resuspended in infection medium and allowed to adhere onto the serum-treated coverslips for 1 hr at 37°C with 5% CO_2_. Neutrophils were treated with M170 anti-CD11b antibody or isotype control, or left untreated, then exposed to CFSE-labeled Ngo. Association and internalization of Ngo by mouse neutrophils was determined as for adherent primary human neutrophils.

### Bead opsonization

Fluoresbrite® Brilliant Blue Carboxylate Microspheres (1.0 µm diameter, Polyscience) were incubated with iC3b (Complement Technology) at 37 °C for 1 hr. Beads were washed and mixed with 3E7 anti-human iC3b antibody (45) (IgG+iC3b opsonized) or an equal volume of PBS (iC3b opsonized) and incubated at 37 °C for 1 hr. Imaging flow cytometry was used to confirm deposition of iC3b ± IgG using FITC anti-C3 and AlexaFluor 647-coupled anti-mouse IgG as in (46).

### Phagosome maturation

Adherent primary human neutrophils were treated with 150 ng mL^−1^ PMA as for experiments with sRBC, then incubated with iC3b- or iC3b + IgG-opsonized beads at a ratio of 1 bead:neutrophil. At the indicated times, the medium was aspirated and neutrophils were fixed in 2% PFA. Neutrophils were stained with FITC-labeled anti-C3 without permeabilization to identify extracellular beads, then permeabilized with ice-cold 1:1 acetone:methanol for 2 min and incubated with rabbit anti-human lactoferrin followed by goat anti-rabbit AlexaFluor 555, or AlexaFluor 555-coupled mouse anti-neutrophil elastase (primary granules).

For Msm, neutrophils were incubated with mCherry-expressing Msm at MOI = 5 for 1 hr. Neutrophils were fixed in 4% PFA, then permeabilized and stained for lactoferrin or elastase as above. OpaD+ and Δopa Ngo were used as positive and negative controls, respectively, for elastase-dependent phagosome maturity.

Intracellular bacteria or beads were considered to be in a positive phagosome if surrounded by > 50% of a ring of fluorescence for the granule protein of interest. 100-200 phagosomes were analyzed per condition. AS and two blinded observers independently quantified the micrographs, and percent positive phagosomes are reported as the average reported by the three observers.

### Talin immunofluorescence

Adherent, IL-8 treated primary human neutrophils were exposed to CFSE-labeled Δopa Ngo as above. After 10 min or 60 min, cells were fixed in 4% PFA. Extracellular Ngo was identified using AlexaFluor 647-coupled anti-Ngo antibody without permeabilization. Cells were re-fixed in PFA, permeabilized with 0.1% saponin in PBS with 10% normal goat serum, and incubated with an Alexa Fluor 555-coupled antibody against talin.

### Complement C3 production by human and mouse neutrophils

Adherent neutrophils (not IL-8 treated) were exposed to 5 µg mL^−1^ cytochalasin B (Sigma) for 20 min followed by 1 µM fMLF (Sigma) for 10 min to stimulate degranulation, or they were instead treated with the DMSO control. Supernatants were passed through a 0.2 µm filter. Δopa Ngo (2 × 10^8^ CFU) was incubated with 1 mL neutrophil supernatant for 20 min at 37° C, pelleted, and washed. Ngo was fixed and processed for imaging flow cytometry, or resuspended in 1x SDS sample buffer for Western blot, in both cases using goat anti-human complement C3 antibody to detect C3 deposition on Ngo. Imaging flow cytometry data acquisition and analysis were performed as previously described for N-CEACAM binding to Ngo (46). Ngo incubated with 10% pooled human serum (Complement Technologies) for 20 min at 37 °C was used as positive control, and Ngo incubated with 10% C3-deficient human serum (Complement Technologies), 10% heat inactivated normal human serum (Sigma), or 10% FBS in RPMI served as negative controls.

### Neutrophil phosphoproteins

Human neutrophils (1 × 10^7^) were suspended in DPBS containing 0.1% dextrose, 1.25 mM CaCl_2_, 0.5 mM MgCl_2_, 0.4 mM MgSO_4_ and incubated with unopsonized Δopa or OpaD+ Ngo for indicated times at 37 °C. Neutrophils were pelleted by centrifugation at 1000 x g at 4°C for 10 min, then resuspended and lysed in 1x SDS sample buffer containing 20 µg/ml aprotinin (ICN), 1 mM PMSF, 20 µg ml^−1^ leupeptin, 20 µg ml^−1^ pepstatin, 40 mM β-glycerophosphate, 10 µM calyculin A, 5 mM NaF, 1 mM Na_3_VO_4_, and 1 mM EDTA. Lysates were separated by SDS-PAGE and analyzed by immunoblot using antibodies for total and phosphorylated AKT, p38, and Erk1/2, followed by goat anti-mouse IgG (H+L) DyLight™ 800 and goat anti-rabbit (H+L) Alex Fluor 680. Bands were quantitated using near-infrared fluorescence detection with the Odyssey imaging system (LI-COR Biosciences).

### Neutrophil degranulation

The surface presentation of CD11a, CD11b (total and active), CD11c, CD63, and CD66b was measured by flow cytometry as in (35). The gate for CD63 or CD66b positivity was set based on the isotype control. Results are reported as geometric mean fluorescence intensity for n=6 biological replicates.

### Reactive oxygen species production

Human neutrophils were exposed to opsonized beads (see above) at a ratio of 100 beads:neutrophil, or exposed to bacteria at MOI = 100. Production of reactive oxygen species over time was measured by luminol-dependent chemiluminescence as in (38). Δopa and OpaD+ Ngo served as negative and positive controls, respectively. Results are representative of at least 3 independently conducted experiments.

### Fluorescence microscopy

Slides were imaged on a Nikon Eclipse E800 UV/visible fluorescence microscope with Hamamatsu Orca-ER digital camera using Openlab (Improvision) or NIS Elements (Nikon) software. Images were exported and processed using Adobe Photoshop CC2019.

### Statistics

Comparisons were made with Student’s *t* test, or one- or two-way ANOVA with appropriate multiple comparisons test as indicated in each figure legend, using Graphpad Prism version 9.2.0 for Windows, GraphPad Software, San Diego, California USA, www.graphpad.com. In experiments with human neutrophils, statistical tests took into account the potential variability of human subjects’ cells by using paired tests. *P* < 0.05 was considered significant, and specific *P* values are indicated in each legend.

## RESULTS

### CR3-dependent phagocytosis elicits weak ROS and primary granule mobilization in human neutrophils

The canonical ligand for CR3 is the iC3b fragment of complement. To define the consequences of CR3 engagement on neutrophil functionality, we exposed fluorescent carboxylate beads that were opsonized with iC3b to adherent, IL-8 treated primary human neutrophils, to mimic the tissue-migrated state of neutrophils responding to infection or injury. First, we confirmed that phagocytosis of iC3b-opsonized particles was CR3-dependent. Adherent, interleukin-8 treated primary human neutrophils were treated with blocking antibodies against CD11b (ICRF44, 44a) or CD18 (TS1/18), or matched isotype controls, then exposed to inert carboxylate beads that were opsonized with iC3b. After 1 hour, cells were fixed and processed for imaging flow cytometry, using an analysis protocol that combines a spot count algorithm and step gate to quantify the binding and internalization of cargo across thousands of cells (43). Coupling of iC3b to the beads was confirmed by imaging flow cytometry **(Fig. S1A-B)**. As expected, association with and internalization of iC3b-opsonized beads by neutrophils was significantly inhibited by CR3 blockade **(Fig. 1A-B)**. Functionality of the CR3 antibodies was confirmed by their ability to inhibit phagocytosis of complement-opsonized sheep erythrocytes **(Fig. S2)**, and CD11b remained on the neutrophil surface after treatment with anti-CD11b antibodies **(Fig. S3A)**.

**Figure 1.**
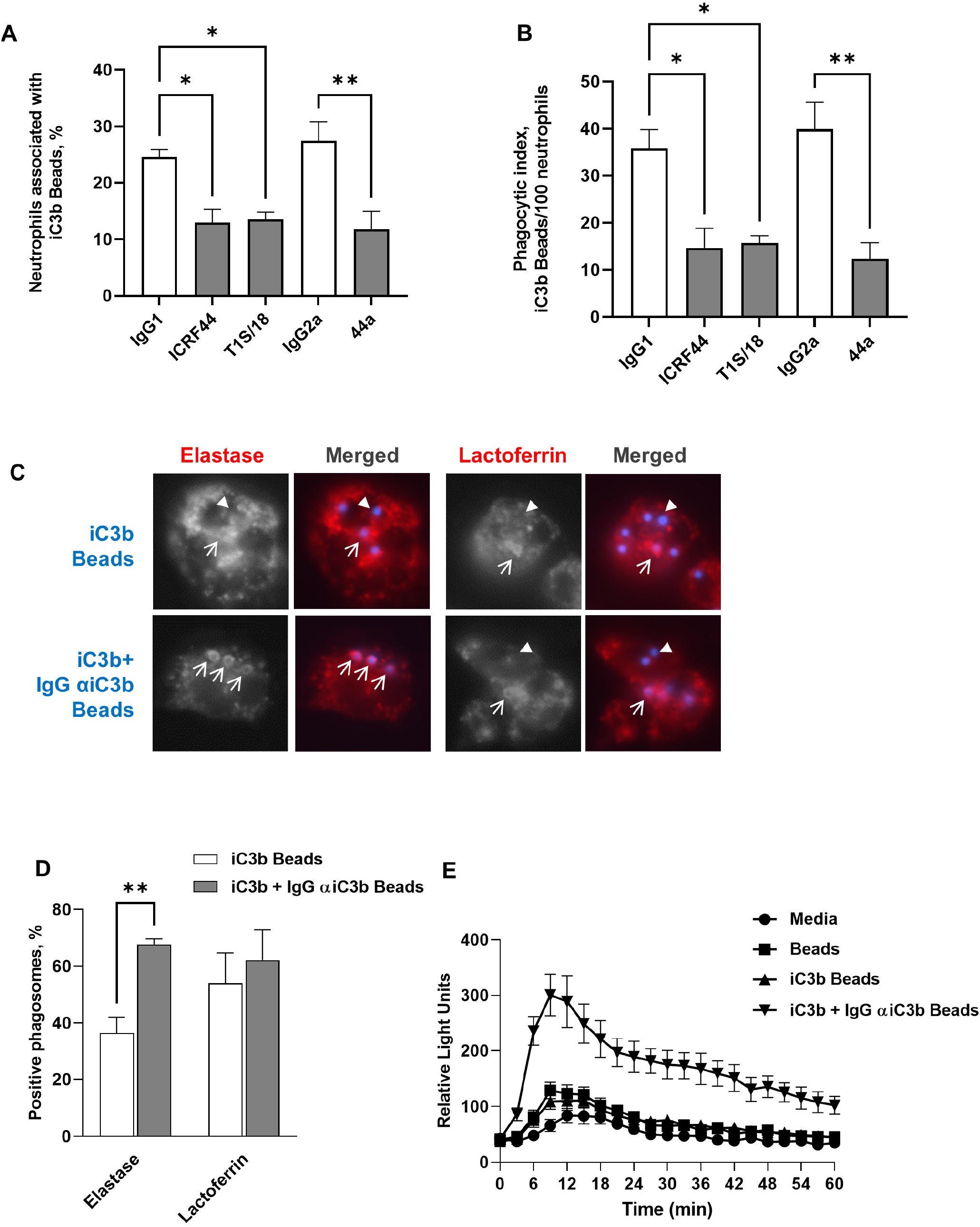
iC3b-coated beads are phagocytosed by neutrophils, but do not stimulate the oxidative burst or full phagosome maturation. **(A-B)** Adherent, PMA-treated primary human neutrophils were exposed to anti-CR3 blocking antibodies (anti-CD11b: ICRF44, 44a; anti-CD18: TS1/18; gray bars) or matched isotype control (white bars), then incubated with Bright Blue™ polystyrene beads opsonized with purified iC3b for 10 min. **(A)** The percent of neutrophils associated with iC3b-opsonized beads was determined by imaging flow cytometry. **(B)** Neutrophils were fixed and processed for immunofluorescence microscopy. The number of phagocytosed iC3b-opsonized beads per 100 neutrophils was determined. For **(A-B)**, data are the mean ± SEM for 3 biological replicates. Statistical significance was determined by ordinary one-way ANOVA with Sidak’s multiple comparisons test. **(C-D)** Adherent, PMA-treated primary human neutrophils were exposed to beads (blue) opsonized with purified iC3b (iC3b) or purified iC3b followed by the 3E7 IgG monoclonal antibody against iC3b (iC3b + IgG αiC3b) for 30 min. Cells were fixed, permeabilized, and stained with an antibody against the primary granule protein neutrophil elastase or the secondary granule protein lactoferrin (red). The red channel is shown in grayscale to better show rings of staining surrounding positive phagosomes (arrows), or their absence around negative phagosomes (arrowheads), alongside the merged images. **(C)** shows representative images. **(D)** The percent of neutrophil elastase- and lactoferrin-positive phagosomes was quantified from over 100 cells from each of 3 biological replicates and presented as the mean ± SEM. Statistical significance was determined by two-way ANOVA with Sidak’s multiple comparisons test. **(E)** Primary human neutrophils were left uninfected (media) or exposed to beads treated with iC3b, iC3b + IgG αiC3b, or untreated. ROS production was measured by luminol-dependent chemiluminescence as the relative light units (RLU) detected at each time point. The graph shown is from one representative experiment of four biological replicates, with three technical replicates per condition. *, *P* ≤ 0.05; **, *P* ≤ 0.01.

The fate of iC3b-opsonized beads and their ability to activate neutrophils were evaluated in comparison to the fate of beads that were opsonized with iC3b, followed by IgG that specifically recognizes human iC3b **(Fig. S1A-B)** (45). iC3b+IgG beads served as a positive control for cargo that stimulates neutrophil ROS production and primary granule release via Fcγ receptors (31). Phagosomes containing iC3b-opsonized beads were significantly less enriched for the primary granule protein neutrophil elastase than beads opsonized with iC3b+IgG **(Fig. 1C-D)**. There was no difference in enrichment of the secondary granule protein lactoferrin on phagosomes between the two bead conditions **(Fig. 1C-D)**. iC3b-opsonized beads induced a minimal ROS response in primary human neutrophils that was identical to what was observed for unopsonized beads, which was less than the response to beads opsonized with iC3b+IgG **(Fig. 1E)**. These findings demonstrate that phagocytosis via CR3 alone weakly activates neutrophil antimicrobial activities.

### CR3 mediates neutrophil phagocytosis of unopsonized Opa-negative *N. gonorrhoeae*

We previously reported that unopsonized, Opa-negative Ngo does not elicit neutrophil ROS and is phagocytosed by adherent neutrophils into primary granule-negative phagosomes, in which they survive(31–33, 47). We tested the hypothesis that Opa-negative Ngo uses CR3 as a phagocytic receptor based on three findings: 1) Complement (serum)-opsonized Ngo also does not elicit ROS from human neutrophils; 2) The percentage of Opa-negative Ngo in immature, elastase-negative phagosomes in adherent human neutrophils is similar to what we observed for serum-opsonized Ngo and for iC3b-opsonized beads **(Fig. 1)** (48); 3) CR3 has been reported as a phagocytic receptor for Ngo in primary cervical epithelial cells (9, 49, 50). To test this hypothesis, adherent, interleukin-8 treated primary human neutrophils were treated with blocking antibodies against CD11b or CD18, or matched isotype controls, and then exposed to piliated Ngo of strain FA1090 in which all *opa* genes were deleted (Δopa) (33). Δopa Ngo was used to limit the potential for variability in results due to Opa phase variation.

Treatment with an antibody against the CD18 α subunit of CR3 significantly reduced both the percent of adherent human neutrophils with intracellular, unopsonized Δopa Ngo **(Fig. 2A)** and the percent of neutrophils with associated (bound and internalized) bacteria **(Fig. 2B)**. Similarly, treatment with an antibody against CD11b (44a (51)) significantly inhibited bacterial association and internalization by adherent human neutrophils **(Fig. 2A-B)**. This result was not specific to Δopa Ngo, as CR3 blocking antibodies also reduced association of neutrophils with predominantly Opa-negative Ngo of the FA1090 RM11.2, MS11 VD300, and 1291 strain backgrounds, in a statistically significant manner **(Fig. S4A)**.

**Figure 2.**
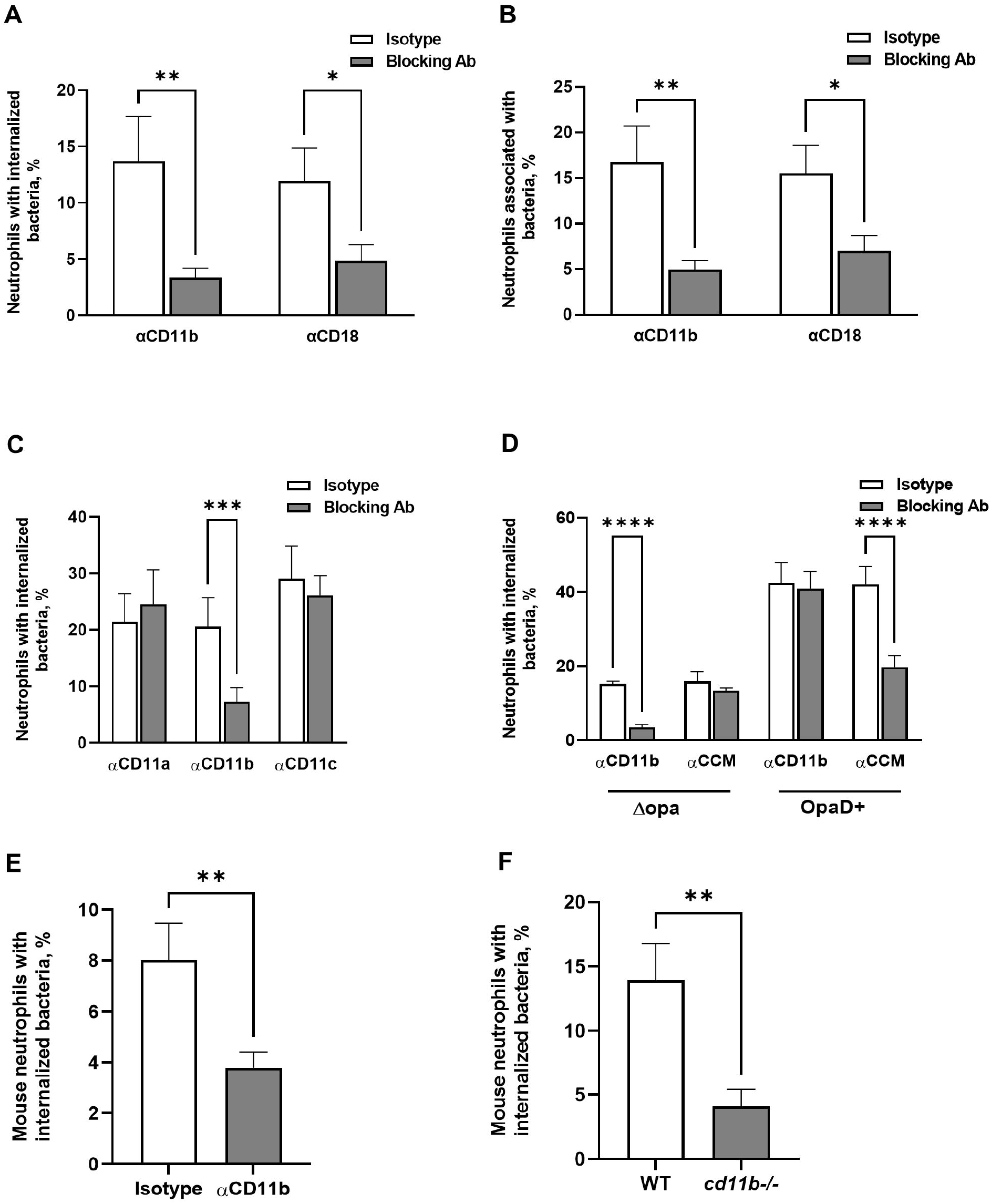
CR3-dependent phagocytosis of Opa-negative *N. gonorrhoeae* by adherent neutrophils. **(A-B)** Adherent, IL-8 treated primary human neutrophils were exposed to blocking antibodies against CD11b (44a) or CD18 (TS1/18) (gray bars), or matched isotype control (white bars). Neutrophils were infected with CFSE-labeled Δopa Ngo for 1 hr, fixed, and stained for extracellular Ngo using a DyLight™ 650 (DL650)-labeled antibody without permeabilization. A step gate was applied to quantify the percentage of neutrophils with internalized **(A)** and associated (bound and internalized) **(B)** Ngo. **(C)** Neutrophils were treated with blocking antibodies against CD11a, CD11b, or CD11c, or isotype control. Internalization of Δopa Ngo was measured as above. **(D)** Neutrophils were treated with blocking antibodies against human CEACAMs (CCM) or CD11b (44a). Internalization of Δopa or OpaD+ Ngo was measured as above. **(E-F)** Neutrophils were purified from the bone marrow of WT or *cd11b-/-* C57BL/6J mice. In **E**, adherent neutrophils from WT mice were exposed to anti-CD11b blocking antibody (M170) or isotype control. Neutrophils were incubated with Tag-IT Violet® labeled Δopa Ngo for 1 hr, fixed, and stained for extracellular bacteria with DL650-labeled anti-Ngo antibody and for the neutrophil surface with FITC-coupled antibody against Ly6G. Cells were analyzed by imaging flow cytometry as above (see **Fig. S5A** for mouse neutrophil gating strategy), and the percent of mouse neutrophils with intracellular bacteria was calculated. Results presented are the mean ± SEM of the following number of biological replicates: (A-B) 4-5; (C), 3-4; (D), 4; (E-F), 6. In **A-D**, statistical significance was determined by two-way ANOVA with Sidak’s multiple comparisons test. In E-F, statistical significance was determined by Student’s paired *t* test. \*\**P* ≤ 0.01, *** *P* ≤ 0.001, **** *P* ≤ 0.0001.

Neutrophil association with and internalization of Δopa Ngo was unaffected by blocking antibodies against CD11a and CD11c **(Fig. 2C, Fig. S4B)**, which dimerize with CD18 and are also expressed on the neutrophil surface **(Fig. S3B)**. Antibody-mediated blockade of human CEACAMs, which are present on the neutrophil surface **(Fig. S3C)** and bind Opa proteins (52), also had no effect on the association and internalization of Δopa Ngo by neutrophils. However, anti-CEACAM antibody inhibited neutrophil association with and phagocytosis of FA1090 Ngo that constitutively expresses the CEACAM1- and CEACAM3-binding OpaD protein (OpaD+) in a statistically significant manner **(Fig. 2D, Fig. S4C)**. CR3-blocking antibodies did not significantly affect the interaction of OpaD+ bacteria with neutrophils, suggesting Opa-CEACAM engagement is dominant over the interaction of Ngo with CR3 **(Fig. 2D, Fig. S4C)**.

Although Ngo is a human-specific pathogen, CR3 is conserved among vertebrates (87.0% similarity and 74.8% identity between *Homo sapiens* (P11215) and *Mus musculus* (E9Q6O4) CD11b). We found that adherent murine neutrophils isolated from bone marrow also bound and phagocytosed Δopa Ngo **(Fig. 2E-F, Fig. S5)**. The association and internalization of Δopa Ngo with mouse neutrophils was significantly reduced upon treatment with anti-CD11b M170 antibody **(Fig. 2E, Fig. S5C)**. Similarly, mouse neutrophils that are genetically deficient for CD11b interacted significantly less with Δopa Ngo than wild-type neutrophils **(Fig. 2F, Fig. S5D)**. Thus CD11b facilitates phagocytosis of adherent Ngo by both human and mouse neutrophils.

The 44a antibody targets a region of human CD11b comprising the I-domain and contributing to the metal ion-dependent adhesion site (MIDAS) motif that is formed upon release of CR3 from an inactive conformation (6, 51). Treatment with the ICRF4 antibody, also directed against the I-domain of active CD11b, significantly reduced Ngo association and internalization by adherent human neutrophils; the M170 antibody, which also blocks the I-domain, particularly the region of CD11b that recognizes complement fragment iC3b, also reduced Ngo-neutrophil interactions, though not in a statistically significant manner (**Fig. S4D-E)**. Treatment of adherent human neutrophils with neutrophil inhibitory factor (NIF), a canine hookworm protein that binds the CD11b I-domain (53, 54), significantly reduced the association and internalization of Δopa Ngo by human neutrophils **(Fig. 3A, Fig. S4F)**. In HL-60 human promyelocytes, which have low levels of surface CD11b, overexpression of full-length CD11b, but not CD11b lacking the I-domain, enhanced phagocytosis of Δopa Ngo, though not in a statistically significant manner **(Fig. 3B-E)**. These results indicate that binding and phagocytosis of Ngo by human neutrophils use the MIDAS and I-domain of CD11b.

**Figure 3.**
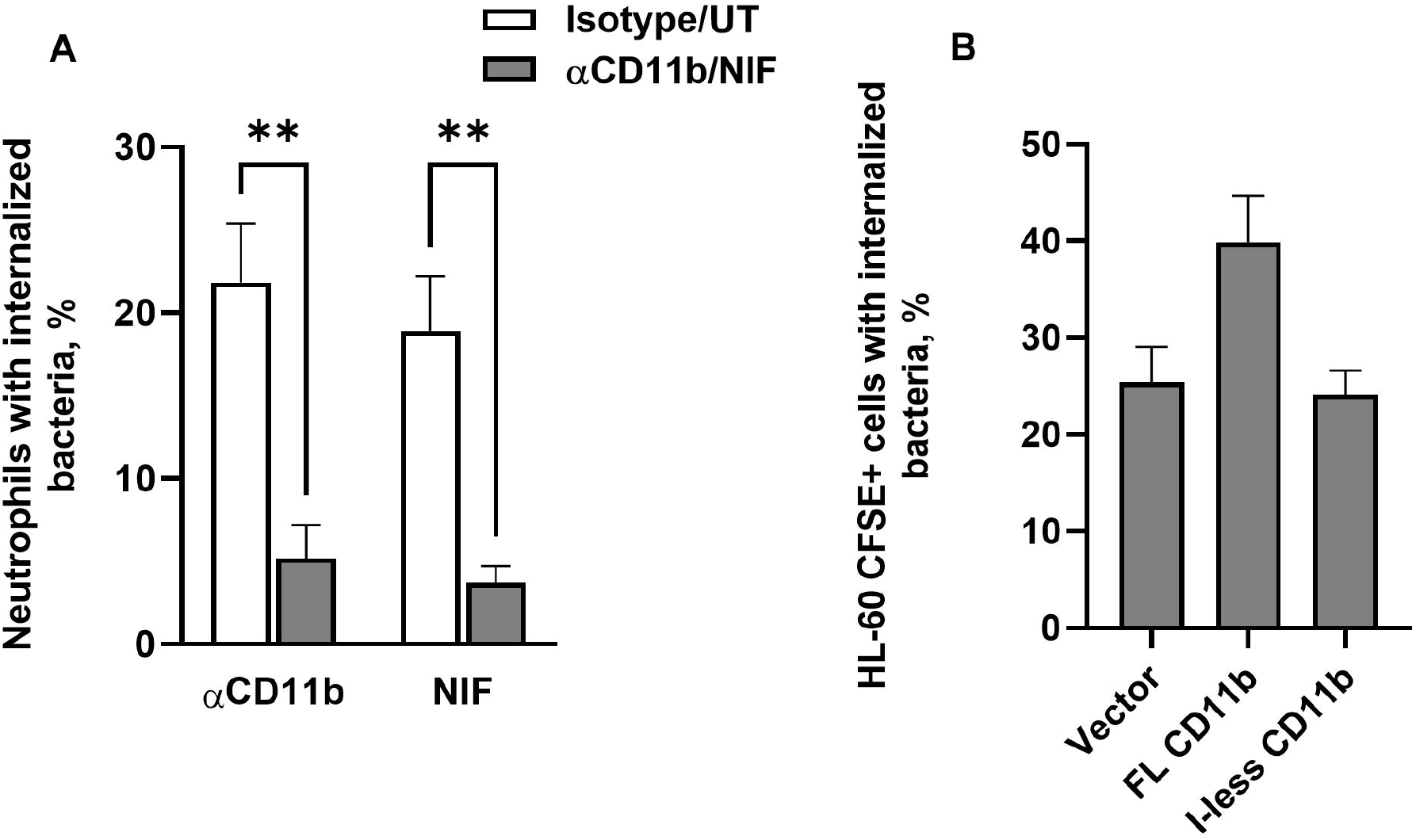
The I-domain and MIDAS motif of CD11b are critical for CR3-mediated phagocytosis of Opa-negative *N. gonorrhoeae*. **(A)** Adherent, IL-8 treated primary human neutrophils were treated with (left group) anti-CD11b (44a; gray bar) or isotype control (white bar), or (right group) *A. caninum* neutrophil inhibitory factor (NIF) (gray bar) or PBS vehicle control (white bar; UT = untreated). The percent of neutrophils with intracellular Ngo was calculated using imaging flow cytometry as in **Fig. 2**. Results are presented as the mean ± SEM for 3 biological replicates, with statistical significance determined by Student’s *t* test for the appropriate pairs. ** *P* ≤ 0.01. **(B)** HL-60 human promyelocytic cells expressing human CD11b that is full-length (FL) or lacking the I-domain (I-less), or carrying empty vector (Vector), were exposed to CFSE-labeled Δopa Ngo for 1 hr. Cells were fixed, stained for extracellular Ngo, and processed for imaging flow cytometry. The percent of CFSE+ HL-60 cells with intracellular Ngo was calculated for 3 biological replicates. Results are presented as the mean ± SEM. Comparisons trended towards but did not reach statistical significance as analyzed using one-way ANOVA.

Neutrophils can make and release complement factors (55–61). However, we did not detect deposition of C3 or C3-derived products on Δopa Ngo that was incubated with primary human or mouse neutrophils **(Fig. S6A, C)**. Ngo incubated with the degranulated supernatant from human and mouse neutrophils also did not show surface C3 reactivity **(Fig. S6B-D)**. We verified the ability to detect complement C3 fragments on Δopa Ngo by opsonizing the bacteria in normal human serum **(Fig. S6B-C)**. Serum opsonization enhanced bacterial phagocytosis by neutrophils, which was significantly reduced by addition of anti-CD11b antibody **(Fig. S6E)**. On cervical cells, Ngo interacts directly with CR3 not only via C3 deposition, but also independently of complement through porin and the glycans on type IV pili (9, 62). CD11b blockade significantly reduced the binding and phagocytosis of the piliated parental Δopa Ngo and an isogenic, hyperpiliated mutant (*ΔpilT*) **(Fig. S7A-B)**. Nonpiliated Ngo (inactivating mutations in the major pilin subunit *pilE* or the pilus secretin *pilQ*) was poorly internalized by neutrophils when compared with the piliated parent, such that any further reduction in internalization by the anti-CD11b antibody was unable to be accurately measured **(Fig. S7A-B)**. Increasing the multiplicity of infection of *ΔpilE* Ngo enhanced bacterial phagocytosis by neutrophils and was significantly inhibited when CD11b was blocked **(Fig. S7C)**. CD11b blocking also significantly reduced the association and internalization of piliated, Δopa Ngo that does not glycosylate pilin, due to mutations in *pglA* or *pglD* **(Fig. S7D-E)**. Thus pili enhance the interaction between Ngo and adherent neutrophils, but phagocytosis of non-piliated Ngo is still inhibited by blocking CR3.

Taken together, these results indicate that phagocytosis of Opa-negative Ngo by adherent neutrophils and neutrophil-like cells uses the CR3 integrin heterodimer, mediated by a region overlapping with the I-domain and MIDAS of CD11b, in a complement-independent manner.

### Activation of CR3 is sufficient to mediate phagocytosis of Opa-negative *N. gonorrhoeae*

Adherent, IL-8 treated neutrophils phagocytose Opa-negative Ngo, albeit less effectively than they phagocytose Opa+ bacteria (30, 31, 33, 48). In contrast, many previous studies reported that neutrophils cannot phagocytose Opa-negative Ngo (21, 23, 28, 63). Notably, these previous studies used neutrophils in suspension and without chemokine treatment, which would keep the cells in a quiescent, inactive state. Thus we hypothesized that Opa-negative Ngo is not phagocytosed by neutrophils in suspension because they have less active surface CR3 than adherent cells. Supporting this hypothesis, primary human neutrophils in suspension had significantly less CD11b on their surface, as well as less CD11b in its active, ligand-binding conformation, compared to adherent, IL-8 treated neutrophils **(Fig. 4A)**.

**Figure 4.**
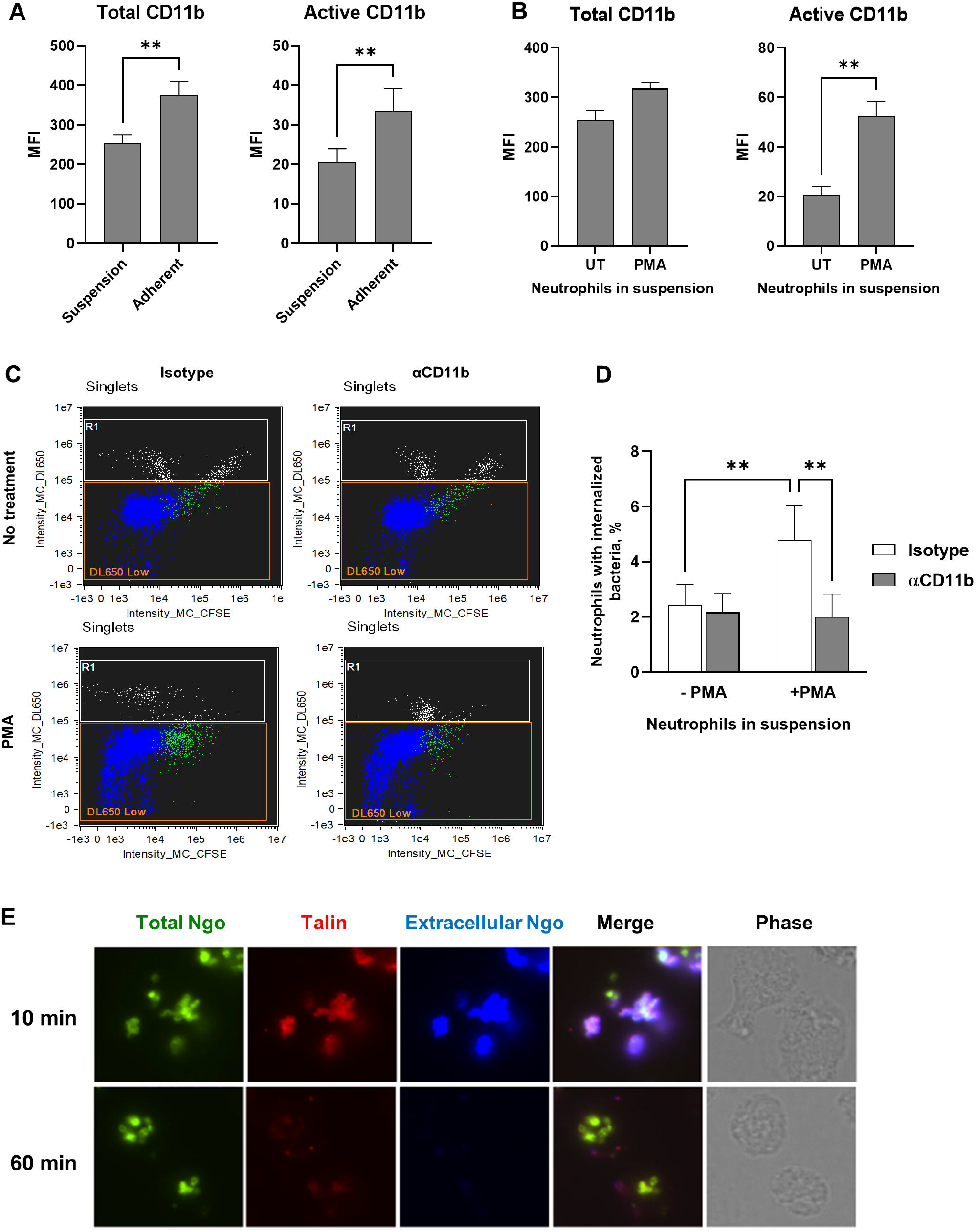
Active CR3 promotes phagocytosis of Opa-negative *N. gonorrhoeae* by human neutrophils in suspension. **(A)** Human neutrophils were maintained in suspension or allowed to adhere to coverslips in the presence of IL-8. The median fluorescence intensity (MFI) of total (left) and active (right) CD11b on the neutrophil surface from n = 5 biological replicates is presented. **(B)** Neutrophils in suspension were left untreated (UT) or treated with PMA, then stained for total (ICRF44; left) and active (CBRM1/5; right) CD11b. The MFI from n= 5 biological replicates is presented. **(C-D)** Untreated or PMA-treated neutrophils in suspension were exposed to anti-CD11b antibody or isotype control, then infected with CFSE-labeled Δopa Ngo at MOI = 1. Phagocytosis was measured by imaging flow cytometry as in **Fig. 2. (C)** Dot plots of the intensity of CFSE (total Ngo) vs. intensity of DL650 (extracellular Ngo). Intact, single neutrophils are identified as the DL650 Low population (blue; R1 = gate for DL650^hi^ cells that are not intact). The green color identifies the population of neutrophils that are associated with CFSE+ Ngo. **(D)** The percentages of untreated and PMA-treated neutrophils with internalized bacteria were calculated as in **Fig. 1**. Results presented are the mean ± SEM from n=4 biological replicates. **(E)** Adherent, IL-8 treated human neutrophils were exposed to CFSE+ Δopa Ngo (green) for 10 min to allow for attachment, or 60 min to allow for internalization. Cells were fixed, stained for extracellular Ngo (blue), then permeabilized and stained for talin (red). Colocalization between extracellular, cell-associated Ngo and talin is seen in the 10 min merged image. Statistical significance was determined by paired Student’s *t* test **(A-B)** or two-way ANOVA followed by Sidak’s multiple comparison test **(D)** for 5 biological replicates. ** *P* ≤ 0.01.

Treatment of suspension neutrophils with phorbol myristate acetate (PMA) significantly increased the amount of surface CD11b in an active conformation, without affecting total surface CD11b **(Fig. 4B)**.

PMA treatment of suspension neutrophils significantly increased their association with **(Fig. S8A)** and internalization of Δopa Ngo **(Fig. 4C-D)**. Addition of CD11b blocking antibody ablated this increase **(Fig. 4C-D, Fig. S8A)**. In contrast, OpaD+ Ngo was bound and internalized by neutrophils in suspension and was unaffected by CD11b blockade **(Fig. S8B-D)**, in keeping with numerous reports that suspension neutrophils phagocytose unopsonized Opa+ Ngo (20, 21, 23, 28, 29, 63–65). These results indicate that phagocytosis of unopsonized, Opa-negative Ngo requires surface-presented CD11b in its active conformation, which is found in adherent and chemokine-primed neutrophils, but not in untreated neutrophils in suspension.

CR3 activation and stabilization of its active high affinity conformation requires the binding of its β integrin tail to talin, which connects the receptor to vinculin and the actin cytoskeleton (6). At 10 min post infection, most Δopa bacteria were surface-bound and colocalized with endogenous talin in adherent, IL-8 treated human neutrophils **(Fig. 4E)**. Colocalization was lost at 60 min post-infection, when Δopa Ngo had been phagocytosed **(Fig. 4E)**. Based on all of these findings, we conclude that adherent, chemokine-treated neutrophils present CR3 in an active conformation that is necessary to mediate binding and phagocytosis of Opa-negative Ngo.

### CR3-dependent phagocytosis is a “silent entry” mechanism of neutrophils

We and others previously reported that Opa-negative Ngo does not stimulate neutrophil ROS production and is found in phagosomes that exhibit delayed fusion with primary granules to enable its intracellular survival (28, 31, 48, 66, 67). To extend these observations, we monitored granule exocytosis and signaling events in adherent, IL-8 treated human neutrophils following exposure to Ngo. There was significantly less surface expression of CD63 on neutrophils that were exposed to Δopa compared with OpaD+ Ngo, indicating a reduction in primary granule exocytosis **(Fig. 5A,C)**. Secondary granule exocytosis, as revealed by CD66b surface exposure, was not significantly different between Δopa and OpaD+ Ngo **(Fig. 5B,C)**. This agrees with our report that Opa- and Opa+ Ngo phagosomes fuse similarly with secondary granules (31). Neutrophils exposed to Δopa Ngo exhibited less phosphorylation of the intracellular signaling kinases Akt, p38, and Erk1/2, compared to those exposed to OpaD+ bacteria at matched time points **(Fig. 5D-I)**. Together, these results connect CR3-mediated phagocytosis of Ngo to a reduced activation state of neutrophils.

**Figure 5.**
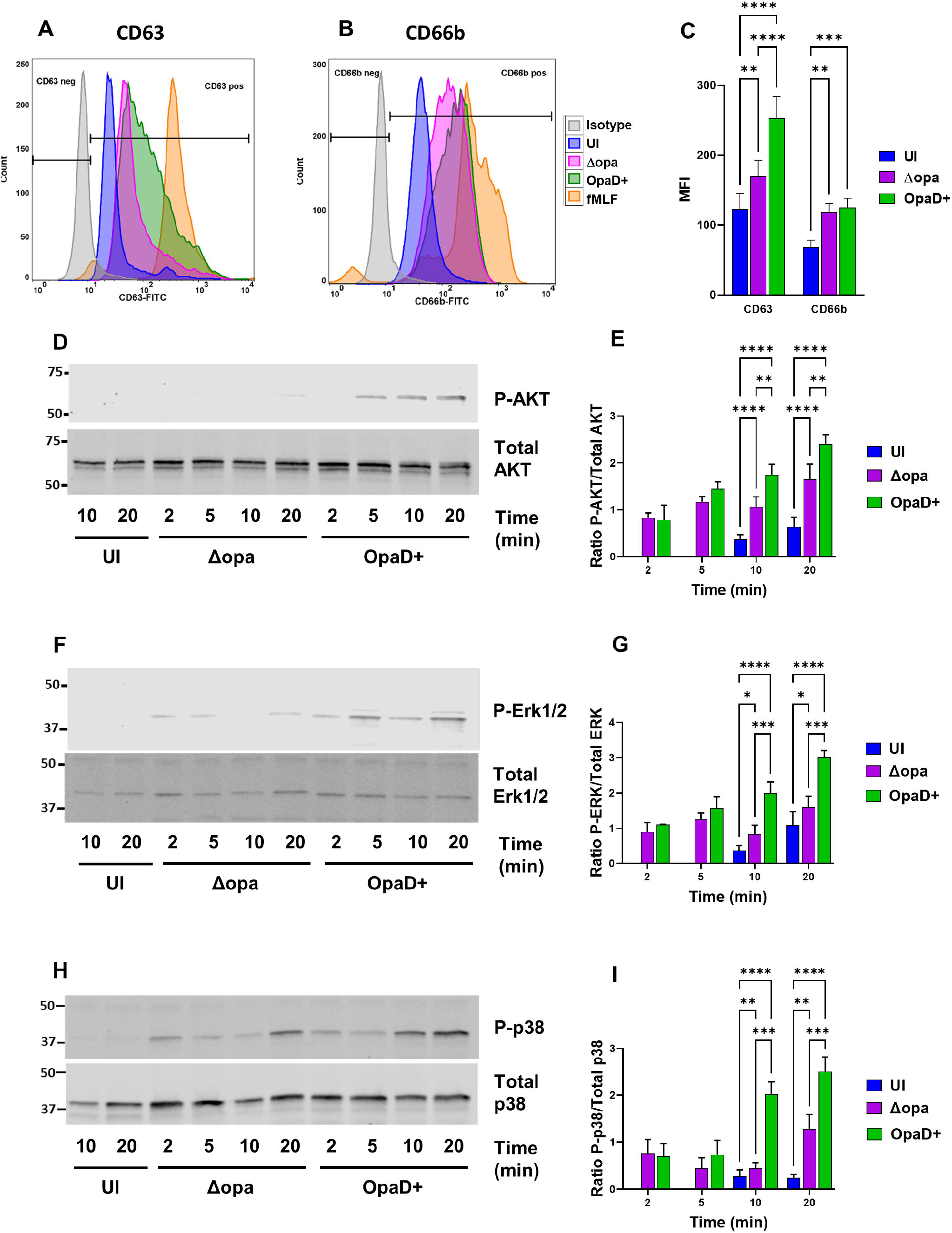
Limited activation of neutrophils following exposure to Opa-negative *N. gonorrhoeae*. **(A-C)** Adherent, interleukin 8-treated primary human neutrophils were exposed to Δopa or OpaD+ Ngo for 1 hr, or left uninfected (UI). Surface presentation of CD63 (primary granules) **(A)** and CD66b (secondary granules) **(B)** was measured by flow cytometry. Neutrophils treated with cytochalasin B + fMLF served as a positive control for degranulation (fMLF). The geometric mean fluorescence intensity (MFI) ± SEM was calculated from *n* = 6 biological replicates **(C)**, with statistical significance determined by two-way ANOVA with Sidak’s multiple comparisons test. **(D-I)** Adherent, IL-8 treated neutrophils were incubated with Δopa or OpaD+ Ngo or left uninfected for the indicated times (UI). Whole cell lysates were immunoblotted for phosphorylated and total p38, ERK1/2, and AKT. **D, F, and H** show a representative blot from one of three biological replicates. **E, G, and I** report the ratio of phosphorylated to total protein by quantitative immunoblot for the three replicates. Statistical significance was determined by two-way ANOVA followed by Tukey’s multiple comparisons test for 3 **(G)** or 4 **(E, I)** biological replicates. * *P* ≤ 0.05, ** *P* ≤ 0.01, *** *P* ≤ 0.001, **** *P* ≤ 0.0001.

To extend these results to an unrelated bacterium, we turned to *Mycobacterium smegmatis*, which is reported to be internalized by neutrophils via CR3 and does not induce primary granule exocytosis (68). We adapted the imaging flow cytometry protocol described above to quantify *M. smegmatis* internalization using a pixel erode mask from the cell boundary **(Fig. S9)**. Phagocytosis of non-opsonized *M. smegmatis* by adherent, IL-8 treated human neutrophils was significantly inhibited when CD18 was blocked **(Fig. 6A)**. Enrichment of neutrophil elastase around *M. smegmatis* phagosomes after 1 hr infection was lower than for Δopa Ngo, and both *M. smegmatis* and Δopa phagosomes were significantly less elastase positive than phagosomes containing OpaD+ Ngo **(Fig. 6B-C)**. Like Δopa Ngo, *M. smegmatis* did not stimulate neutrophil ROS production, in contrast to the strong ROS response elicited by OpaD+ Ngo **(Fig. 6D)** (33, 47).

**Figure 6.**
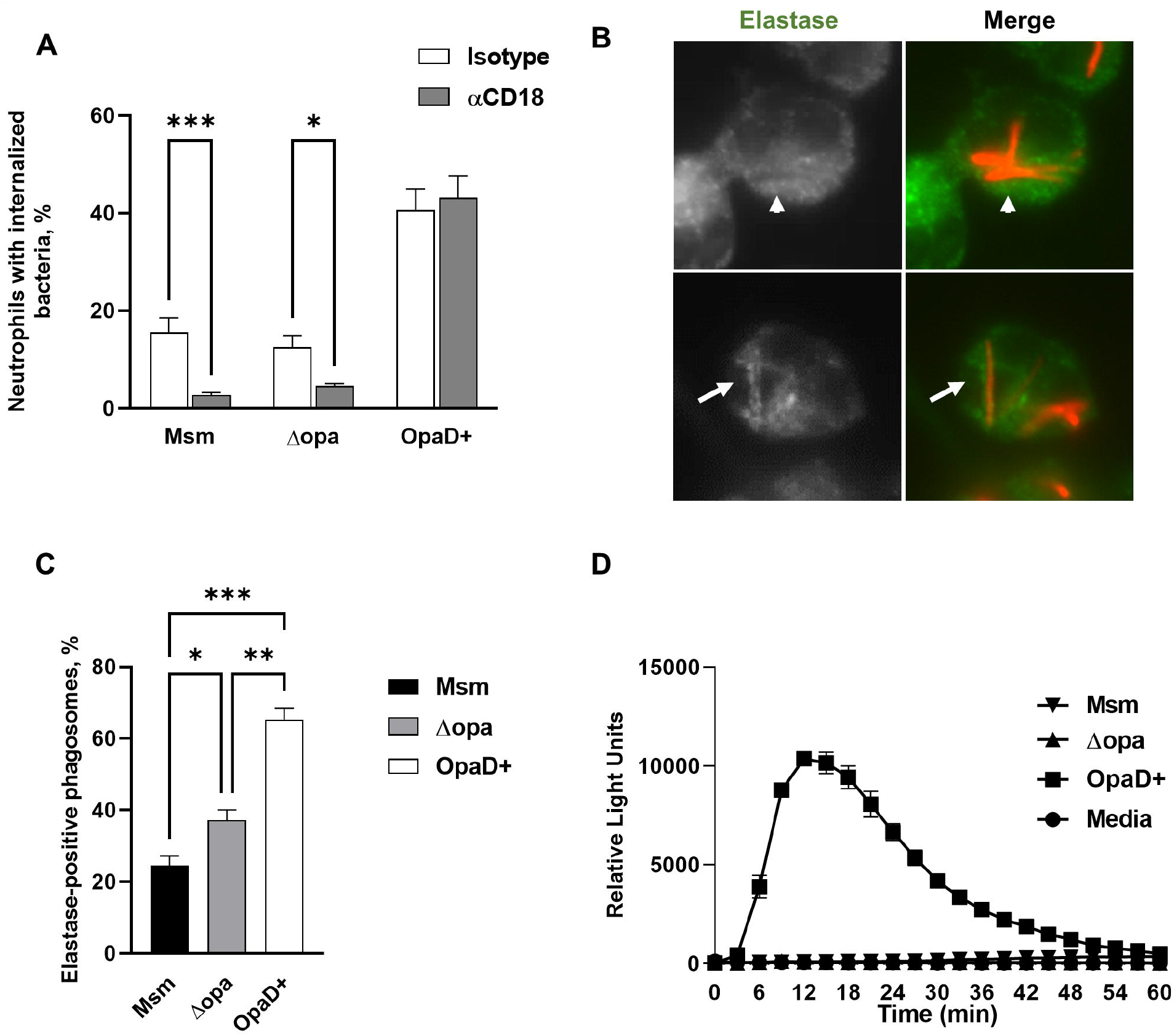
Phagocytosis of *M. smegmatis* by human neutrophils is CR3-dependent and does not stimulate the oxidative burst or full phagosome maturation. **(A)** Adherent, IL-8 treated human neutrophils were treated with either isotype control or anti-CD18 antibody, then exposed to Tag-IT Violet™ labeled *M. smegmatis* (Msm) for 1 hr. Neutrophils were fixed, stained for CD11b, and examined by imaging flow cytometry. Intracellular bacteria were defined as those within a mask set by the CD11b surface staining eroded by 4 pixels and quantified by spot count (see **Fig. S9**). The percent of neutrophils with intracellular *M. smegmatis*, Δopa Ngo, and OpaD+ Ngo, with and without CD18 blockade, was calculated from 5 biological replicates and presented as the mean ± SEM. **(B-C)** Adherent, IL-8 treated human neutrophils were infected with mCherry-expressing *M. smegmatis* at MOI = 5 for 1 hr (red). Cells were fixed, permeabilized and stained for neutrophil elastase (green). Two representative neutrophils with *M. smegmatis* are shown in **(B)**, with the grayscale elastase signal on the left and merged fluorescence image with *M. smegmatis* on the right. The arrow identifies an elastase-positive phagosome, and the arrowhead identifies an elastase-negative phagosome. The percent of elastase-positive phagosomes containing *M. smegmatis*, Δopa Ngo, and OpaD+ Ngo was quantified from 50-100 phagosomes per condition. Results presented are the mean ± SEM from 4 biological replicates. **(D)** Primary human neutrophils were left uninfected (media) or exposed to *M. smegma*tis, or Δopa or OpaD+ Ngo at MOI = 100. ROS production by luminol-dependent chemiluminescence was measured as in **Fig. 1E**. The graph shown is from one representative of 3 biological replicates. Each data point is the mean ± SEM from three technical replicates. Statistical significance was determined by two-way ANOVA followed by Sidak’s multiple comparisons test **(A)** or repeated measures one-way ANOVA followed by Tukey’s multiple comparisons test. * *P* ≤ 0.05, ** *P* ≤ 0.01, *** *P* ≤ 0.001.

Together, the results with Ngo, iC3b-opsonized beads, and *M. smegmatis* support a model in which CR3 engagement facilitates phagocytosis of diverse cargo with incomplete neutrophil activation, and this is exploited by pathogens to evade full neutrophil antimicrobial activity.

## DISCUSSION

CR3 is a prominent phagocytic receptor on vertebrate phagocytes, including neutrophils. Despite extensive research on how CR3 is activated to mediate phagocytosis, the downstream consequences of CR3-mediated phagocytosis remain uncertain. Here, we use iC3b-opsonized polystyrene beads to show that phagocytosis solely by CR3 in neutrophils does not elicit a prominent ROS response or primary granule release. The same phenotypes are found for neutrophil CR3-binding *M. smegmatis*. For the first time, we identify CR3 as the main phagocytic receptor on neutrophils for Ngo that is not opsonized or lacks other adhesins (e.g. phase variable Opa proteins). Phagocytosis of Ngo requires the I-domain and MIDAS of CD11b, and placement of CR3 into an active conformation that is only achieved in adherent, primed neutrophils, and not for unstimulated neutrophils in suspension. CR3-mediated phagocytosis of Opa-negative Ngo was linked to a less activated state of neutrophils, with reduced activation of protein kinases implicated in proinflammatory signaling, less granule fusion with the cell surface (less degranulation) and phagosomes (less phagosome maturation), and minimal ROS production. These results with three unrelated phagocytic cargo define CR3-mediated phagocytosis as a pathway of “silent entry” into neutrophils, which is exploited by pathogens like Ngo and Msm to create an intracellular niche that supports their survival **(Fig. 7)**.

**Figure 7.** CR3-mediated phagocytosis is a silent entry pathway into neutrophils. Non-opsonized substrates that are phagocytosed into neutrophils via CR3 do not induce the oxidative burst, poorly activate inflammatory signaling cascades, and predominantly reside in immature phagosomes. While this pathway of silent entry would facilitate non-inflammatory clearance of iC3b-tagged apoptotic cells and immune complexes (recapitulated in this study by use of iC3b-coated polystyrene beads), it is exploited by diverse microbes, including *Neisseria gonorrhoeae* and *Mycobacterium smegmatis*, to avoid degradation and killing within neutrophils.

Despite decades of study on CR3, there are conflicting results regarding the downstream fate of cargo that engage CR3 on phagocytes. iC3b-opsonized zymosan, a common experimental substrate for neutrophils, induces a potent ROS and degranulation response (69). However, zymosan is also recognized by dectin receptors that activate neutrophils (70, 71). The β-glucan of zymosan also interacts directly with the lectin domain of CD11b (69). Erythrocytes coated with either C3b or iC3b are phagocytosed but do not induce ROS generation in human monocytes and neutrophils (72). In contrast, C3 and iC3b-coated latex beads are reported to elicit neutrophil ROS (73). Here, we found that iC3b-opsonized polystyrene beads, in the absence of any other ligands, are phagocytosed by neutrophils into immature phagosomes and do not induce ROS production. Given the results with zymosan, we suggest that inconsistencies in the literature regarding CR3-mediated phagocytosis and neutrophil activation are attributable to other surface features of the phagocytosed target, or the presence of other soluble components that opsonize the target or stimulate neutrophils. For instance, binding of C5a, the cleavage product of complement component 5, to the C5a receptor on neutrophils leads to upregulation of CR3 on neutrophils, and consequent ROS production (74–76). It is intriguing that pathogen-associated molecular patterns on Δopa Ngo do not activate neutrophils in the infection condition used here, although Ngo lipooligosaccharide is a ligand for TLR4 and its PorB porin binds TLR2 (77, 78). In macrophages, engagement of CR3 by complement-opsonized *Francisella tularensis* downregulates TLR2 responses to dampen proinflammatory signaling (79). If this is a general approach used by microbes that target CR3, then bacteria as unrelated as Ngo and Msm would be expected to exploit CR3 not just to access an intracellular niche in phagocytes that lacks full degradative capacity, but also to lessen the host-protective inflammatory response.

How Ngo interacts with CR3 on neutrophils is not currently known. We did not detect any deposition of C3 on Ngo that was incubated with neutrophils or degranulated neutrophil supernatant **(Fig. S6)**. This is in contrast to the action of primary human cervical cells, which release C3 that covalently reacts with Ngo lipooligosaccharide (80). Ngo also did not use its pili to mediate CR3 binding and phagocytosis by adherent neutrophils **(Fig. S7)**. Here, pili may be dispensable because CR3 is already in its active conformation on adherent, IL-8 treated neutrophils, whereas in cervical cells, the interaction of glycosylated pili with CR3 is required to activate CR3 (62). The interaction between cervical CR3 and Ngo involves a cooperative synergy with pili and porin (50). However, we were unable to test a role for porin in CR3-mediated phagocytosis by neutrophils because porin is essential in Ngo. The monoclonal antibody that recognizes FA1090 porin cannot be processed into F’(ab) fragments for blocking, and the intact antibody would opsonize the bacteria for phagocytosis via Fc receptors. We envision three possibilities to explain how Δopa Ngo uses CR3 on neutrophils for phagocytosis. First, Ngo surface structures other than pili and porin directly engage CR3. For instance, in the related pathogen *Neisseria meningitidis*, lipooligosaccharide interacts with CR3 (81). Second, Ngo binds a neutrophil-derived protein such as LL-37 or myeloperoxidase, which have been reported as ligands for CR3 (13, 82). Third, CR3 signaling may be required for activation of another receptor that drives Ngo phagocytosis by neutrophils, or for stimulating actin-dependent membrane ruffling and bacterial uptake by macropinocytosis. In all cases, the consequence would be phagocytosis without ROS production or full degranulation, and ineffective bacterial killing.

While neutrophils in suspension cannot phagocytose unopsonized, Opa-negative Ngo, neutrophils that are adherent and treated with IL-8 can (28-31, 33, 63, 83). These discordant results can be explained by our finding that neutrophils in suspension have reduced levels of CR3 on their surface, including CR3 in its active ligand-binding conformation, compared to adherent and primed neutrophils. In circulation, CR3 is inactive to prevent indiscriminate binding of phagocytes to endothelial cells. Both inside-out and outside-in signals, including attachment to extracellular matrix proteins such as fibronectin, activate CR3 on phagocytes (5, 6). We found that treating neutrophils in suspension with PMA, a PKC activator known to promote CR3 activation (84), was sufficient to increase phagocytosis of Δopa Ngo in a CR3-dependent manner **(Fig. 3)**. This result highlights the importance of the physiological state of immune cells in host-pathogen interactions. Notably, adherent neutrophils required PMA treatment in order to phagocytose iC3b-coated beads or erythrocytes, but they successfully phagocytosed Ngo and *M. smegmatis*. This result may indicate that a second signal, such as a microbe-associated molecular pattern on the bacterial surface, is required for inside-out signaling to place neutrophil CR3 in an active phagocytosis-competent state. Alternatively, Ngo and *M. smegmatis* may interact with a slightly different domain of CR3 than iC3b. This possibility is supported by the differential effects of the anti-CD11b blocking antibodies 44a and M170 on neutrophil phagocytosis of Δopa Ngo, compared with iC3b-opsonized sRBCs **(Fig. 2, Fig. S4)**. M170 had a statistically significant effect on phagocytosis by mouse neutrophils compared with human neutrophils, which could be due to a higher affinity of this antibody for mouse vs. human CD11b, or differences in the activation state of CR3 in mouse vs. human neutrophils in our experimental conditions. We noted that anti-CEACAM antibody reduced phagocytosis of OpaD+ Ngo by neutrophils to what was measured for Δopa Ngo without CR3 blocking **(Fig. 2D)**. This result implies that CR3 and CEACAM independently drive neutrophil phagocytosis of Opa-negative or Opa+ Ngo, respectively. This scenario differs from IgG receptor signaling, where CR3 synergizes with Fcγ receptors to promote rapid phagocytosis and degradation of substrates (85). The dynamics between CR3 and other phagocytic receptors, particularly CEACAMs, in neutrophil phagocytosis, and how additional cargo-derived signals affect phagocytosis, warrants further investigation. However, these questions will require a more genetically tractable system than primary human neutrophils, which are terminally differentiated.

Diverse pathogens, as well as their secreted toxins and virulence factors, interact with CR3 in a complement-independent manner (7, 8, 14, 68, 79, 81, 86–91). This study directly connects engagement of CR3 to evasion of phagocyte activation and phagocytic clearance, which we are terming “silent entry.” Furthermore, it identifies CR3 as a participant in a pathway that is exploited by *M. smegmatis* and Ngo to be taken up by and survive inside neutrophils. This may help explain the clinical observation that neutrophils in human gonorrheal exudates contain intact intracellular Ngo (19). Our work is in agreement with reports that the phagocytosis of *Mycobacterium smegmatis, M. kansasii*, and *M. tuberculosis* by neutrophils or macrophages uses CR3 and does not lead to ROS production or phagosome maturation (14, 68, 86, 92–96). Similarly, *Leishmania* prevents ROS production and phagolysosome formation in phagocytes in a CR3-dependent manner to enable its survival (97, 98), and macrophages infected with complement-opsonized *F. tularensis* have limited pro-inflammatory signaling and cytokine production (79, 99). However, our work with OpaD+ Ngo suggests that the exploitation of CR3 by pathogens can be overcome by ectopically activating ITAM-bearing receptors such as Fcγ receptors, dectins, and CEACAM3, which send signals that activate phagocytes alongside or independently of CR3 (6, 85, 100, 101). Vaccination or host-directed therapies can therefore be used to target the phagocytes that are recruited to sites of infection in order to thwart this silent entry pathway and therefore enhance pathogen clearance.

## Supporting information

All supplemental figures and legends

## ACKNOWLEDGEMENTS

We thank Michael Solga and Joanne Lannigan (University of Virginia Flow Cytometry Core, RRID: SCR_017829) for their contributions to the imaging flow cytometry protocols used in this study. We thank Michael Apicella, Jennifer Edwards, Girija Ramakrishnan, Hank Seifert, and Christina Stallings for strains and mutant constructs, Victor Torres for HL-60 cell lines, and Larry Schlesinger for helpful discussions. We thank CJ Anderson, Borna Mehrad, and Kathy Michaels for advice on isolation and immunofluorescence of mouse neutrophils, and Jeff Teoh for assistance with establishing flow cytometry protocols. We thank Marieke Jones for advice on statistical analyses and Allison Alcott for assistance with manual quantitation. This work was supported by NIH R01 AI097312, the University of Virginia School of Medicine Pinn Scholar Award, and the Department of Microbiology, Immunology, and Cancer Biology to AKC. SAR was supported in part by NIH T32 AI007046. SAR and LMW were supported in part by the Robert R. Wagner Fellowship, University of Virginia. KPD was supported in part by the University of Virginia’s Harrison Undergraduate Research Fellowship and Hutcheson Family Fellowship. Human subjects gave informed consent and participated in research according to a protocol approved by the University of Virginia Institutional Review Board for Health Sciences Research. Research with animals was conducted according to a protocol approved by the University of Virginia Animal Care and Use Committee.

## Graphical Abstract

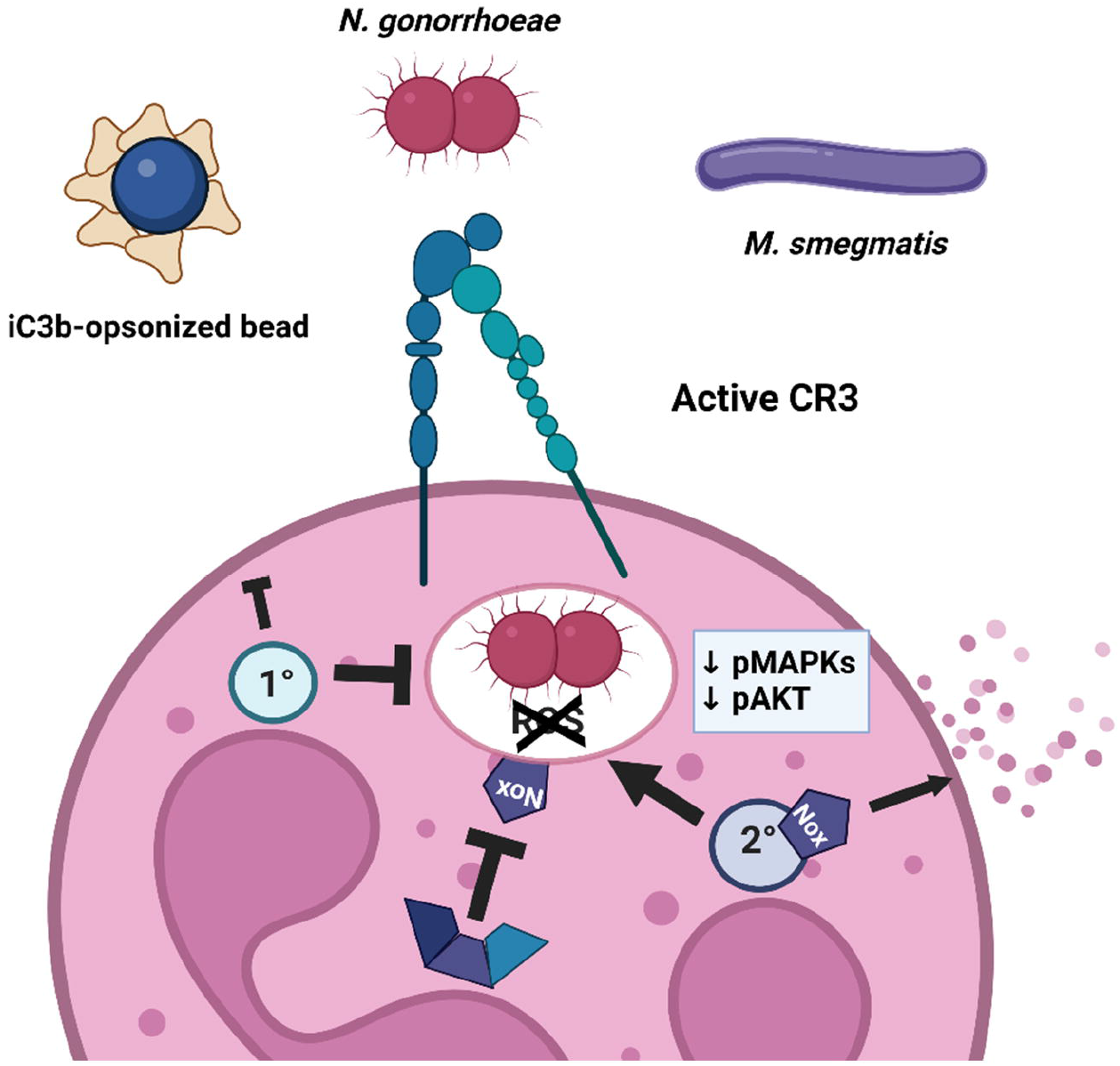

