## Supplementary material for "Phagocytosis via complement receptor 3 enables microbes to evade killing by neutrophils": All supplemental figures and legends

**Legends for Supplemental Figures**

**Figure S1. iC3b and anti-iC3b are present on the surface of opsonized polystyrene beads.** Brilliant Blue ™ polystyrene beads were left unopsonized (Beads), opsonized with human iC3b (iC3b beads), or opsonized with iC3b followed by 3E7 monoclonal antibody against human iC3b (iC3b + IgG αiC3b beads). Beads were stained with FITC-coupled antibody against human C3 and Alexa Fluor 647-coupled antibody against mouse IgG, then analyzed by imaging flow cytometry. **(A)** Flow cytometry plots of FITC vs. AF647 with the percent of negative, single positive, and double positive beads shown. **(B)** Representative images from unopsonized beads (top panel), iC3b-opsonized beads (middle panel), and iC3b-anti-iC3b opsonized beads (lower panel).

**Figure S2. Anti-CR3 antibodies block phagocytosis of serum opsonized sheep red blood cells (RBCs) by primary adherent IL-8 primed human neutrophils**. **(A,B)** Adherent primary human neutrophils were treated with 150 ng $mL^{-1}$ phorbol 12-myristate 13-acetate (PMA), followed by the indicated blocking antibodies against CD11b (ICRF44, 44a) or CD18 (TS1/18) (gray bars), or matched isotype controls (white bars). Neutrophils were then incubated for 10 min with sheep RBCs that were opsonized in 10% human serum. Extracellular RBCs were lysed with water, cells were fixed, and phase contrast images were captured for 10 randomly selected fields of view. **(A)** Representative phase contrast images of neutrophils treated with isotype control (upper panel) or 44a anti-CD11b antibody (lower panel). Arrows indicate internalized RBCs, arrow heads point to membranes of lysed extracellular RBCs. Scale bar, 10 µm. **(B)** The phagocytic index was calculated as the number of internalized RBCs per 100 neutrophils. Values are mean ± SEM for 3-12 biological replicates. Statistical significance was determined by one-way ANOVA with Sidak's multiple comparisons test (*, $P \leq 0.05$; ***, $P \leq 0.001$). **(C)** Serum-opsonized sheep RBCs (opsRBCs; left) and non-opsonized RBCs (non-opsRBCs; right) were processed for immunofluorescence with a FITC-coupled antibody against human C3, to visualize C3 deposition specifically on the opsonized RBCs. Scale bar, 10 µm.

**Figure S3. CD11 isoforms and CEACAMs are present at the surface of adherent, IL-8 treated primary human neutrophils and are bound by blocking antibodies**. Adherent, IL-8 treated primary human neutrophils were exposed to 20 µg $mL^{-1}$ of the indicated antibodies or isotype control for 1 hr, stained with anti-mouse secondary antibodies, and analyzed by flow cytometry. Results are presented as representative histograms of cell counts (y axis) per fluorescence intensity measurement (x axis). **(A)** Antibodies against CD11b (M170 and 44a). **(B)** Antibodies against CD11a, CD11b (44a), and CD11c. **(C)** Antibodies against CEACAM (CCM) and CD11b (44a).

**Figure S4. The association of multiple strains of Opa-negative *N. gonorrhoeae* with neutrophils is CR3-dependent. (A)** Adherent, IL-8 treated primary human neutrophils were treated with 44a anti-CD11b antibody (gray bars) or isotype control (white bars), then infected with CFSE-labeled piliated, Opa-negative Ngo of the following strain backgrounds: FA1090 1-81-S2 and RM11.2, MS11 VD300, and 1291. FA1090 Δopa (see **Fig. 2**) was included for comparison. The percent of neutrophils that are positive for associated Ngo (focused singlets, $DL650^{low}$ $CFSE^{+}$) was measured by imaging flow cytometry as described in **Fig. 2**. **(B)** Neutrophils were treated with blocking antibodies against CD11a, CD11b (44a), or CD11c, or isotype control, then exposed to Δopa Ngo. The percent of neutrophils that are positive for associated Ngo was measured as above. **(C)** Neutrophils were treated with blocking antibodies against human CEACAMs (CCM) or CD11b (44a), or isotype controls, then exposed to Ngo. The percent of neutrophils that are positive for associated Δopa or OpaD+ Ngo was measured as above. **(D-E)** Neutrophils were treated with ICRF44 and M170 blocking antibodies against CD11b, or matched isotype control, then exposed to Δopa Ngo. The percent of neutrophils positive for associated **(D)** or internalized **(E)** Ngo was measured as above. **(F)** Neutrophils were treated with neutrophil inhibitory factor (NIF) from *Ancylostoma caninum*) (gray bar) or left untreated (UT; white bar). In parallel, neutrophils were treated with anti-CD11b (44a; gray bar) or isotype control (white bar). The percent of neutrophils that are positive for associated Δopa Ngo was measured as above. Data presented are the mean ± SEM for the following number of biological replicates: **(A)** 4-22; **(B)** 4; **(C)** 4; **(D)**, 3. Statistical comparisons were made using two-way ANOVA followed by Sidak's multiple comparisons test. ** $P \leq 0.01$, *** $P \leq 0.001$, **** $P \leq 0.0001$.

**Figure S5. Adherent mouse neutrophils associate with Opa-negative *N. gonorrhoeae* in a CD11b-dependent manner.** Neutrophils that were purified from bone marrow of WT or *cd11b-/-* C57BL/6J mice were allowed to settle on serum-coated coverslips, then exposed to Tag-IT Violet™ labeled Δopa Ngo for 1 hr. Cells were fixed, stained with anti-Ly6G-FITC and anti-Ngo DL650 antibody, and analyzed by imaging flow cytometry. **(A)** Gating strategy: i) Single cells were obtained by gating on objects with low area and high aspect ratio values. ii) Focused cells were identified by gating on objects with high gradient RMS. iii) Intact cells were defined as a population of cells with low DL650 intensity. iv) Neutrophils were identified as Ly6G+ cells. v) Spot count feature was used to identify neutrophils not associated with bacteria (No bacteria) and associated with bacteria (cells with ≥1 Tag-IT Violet™ positive spots). vi) For neutrophils with 1-3 Tag-It Violet™ positive spots, the spot count feature was used to identify and count the number of DL650+ spots. vii) A step gate was applied to identify and count the number of neutrophils with at least 1 Tag-IT Violet™ positive, DL650-negative (intracellular) bacteria. **(B)** Representative images of Ly6G+ mouse neutrophils with internalized bacteria (top panel), and both internalized and surface bound bacteria (middle and bottom panel) examined by imaging flow cytometry. **(C)** Adherent mouse neutrophils were treated with anti-CD11b (M170; gray bar) or isotype control (white bar), then infected with Δopa Ngo. The percent of neutrophils with associated bacteria was calculated as above. **(D)** Adherent neutrophils from wild-type (white bar) or isogenic *cd11b-/-* (gray bar) mice were infected with Δopa Ngo, and the percent of neutrophils with associated Ngo was calculated as above. Data are the mean ± SEM from 6 biological replicates. Statistical significance was determined by paired Student's *t* test. ** $P \leq 0.01$.

**Fig. S6. Human neutrophils are not a source of C3 for opsonization of *N. gonorrhoeae.* (A)** Adherent, IL-8 treated primary human neutrophils were exposed to non-opsonized Δopa Ngo for 15 min, fixed, and stained with FITC-labeled anti-C3 (green) and DL650-labled anti-Ngo (red) antibodies. Fluorescence micrographs were captured for the FITC and DL650 channels (presented as grayscale), with the merge in color showing overlap with the phase image for the neutrophils. **(B)** Δopa Ngo was incubated with normal human serum as a positive control for C3 opsonization (third panel), or with the supernatants collected from adherent primary human neutrophils that were left untreated (first panel) or treated with cytochalasin B (cytoB) and fMLF to stimulate degranulation (second panel). Bacteria were washed and stained with FITC-labeled anti-C3 antibody. Representative fluorescence micrographs with the FITC channel overlaid on the phase image are presented. **(C)** Neutrophils were treated with cytoB + fMLF as above, or left untreated (-). Neutrophils were exposed for 15 min to unopsonized Δopa Ngo. Neutrophils were fixed, stained with FITC-labeled anti-C3 antibody and DL650-labeled anti-Ngo antibodies, and analyzed by imaging flow cytometry. The spot count feature was used to quantify cells with positive FITC signals within the population of neutrophils containing 1-3 DL650+ Ngo for 3 biological replicates. Untreated neutrophils exposed to Δopa Ngo that was opsonized in normal human serum (ops) served as a positive control. **(D)** Δopa Ngo was left untreated, or incubated with the supernatant from adherent mouse neutrophils that were untreated (Mouse Neutrophil Supt) or treated with cytoB + fMLF (+fMLF Mouse Neutrophil Supt). Ngo was incubated with 50% normal mouse serum as a positive control (+NMS) or PBS as a negative control (-). Ngo was washed and stained with FITC-labeled anti-mouse C3 antibody and analyzed by imaging flow cytometry as in **C**. The percent of bacteria that were in the C3+ gate was quantified from 3 biological replicates. **(E)** Adherent, IL-8 treated primary human neutrophils were exposed to anti-CD11b (44a; gray bars) or isotype control (white bars), then infected with Δopa Ngo that was not opsonized (left) or opsonized in normal human serum as in **B** (right). The percent of neutrophils with internalized Ngo was quantified by imaging flow cytometry as in **Fig. 2**. Results presented are for 4-6 biological replicates. Statistical significance in **C-D** was determined by ordinary one-way ANOVA with Tukey's multiple comparisons test, and in **E** by two-way ANOVA with Sidak's multiple comparisons test. * $P \leq 0.05$,** $P \leq 0.01$***, $P \leq 0.001$, **** $P \leq 0.0001$.

**Figure S7. Piliation enhances association and internalization of Opa-negative *N. gonorrhoeae* with primary human neutrophils, and phagocytosis of nonpiliated bacteria is blocked with anti-CR3 antibody. (A,B)** Adherent, IL-8 treated primary human neutrophils were exposed to anti-CD11b antibody (44a; gray bars) or isotype control (white bars). Neutrophils were then exposed to the piliated Δopa Ngo parent, isogenic non-piliated mutant Ngo (*ΔpilE, ΔpilQ*), or isogenic hyperpiliated, retraction-deficient *ΔpilT* Ngo, all labeled with CFSE, for 1 hr. The percent of neutrophils with internalized **(A)** and associated **(B)** Ngo was determined using imaging flow cytometry as in **Fig. 2**. Data are the mean ± SEM for 7-17 biological replicates. **(C)** Adherent neutrophils were exposed to CFSE-labeled piliated, Δopa Ngo or isogenic *ΔpilE* mutant at low (MOI=1) and high (MOI=50) MOI. Internalization was defined as CFSE fluorescence within a mask created from the phase image, eroded by 4 pixels from the cell surface. The normalized intracellular CFSE was measured for 3 biological replicates. **(D, E)** Adherent human neutrophils were exposed to piliated Δopa Ngo, or isogenic pilin glycosylation-deficient *ΔpglA* and *ΔpglD* bacteria, which were labeled with CFSE. The percent of neutrophils with associated **(D)** and internalized **(E)** Ngo was quantified by imaging flow cytometry as above. Results are the mean ± SEM for 4-6 biological replicates. Statistical significance was determined by two-way ANOVA followed by **(A-C)**

Tukey's multiple comparisons test or **(D-E)** Sidak's multiple comparisons test. ** $P \leq 0.01$***, $P \leq 0.001$, **** $P \leq 0.0001$.

**Figure S8. Active CR3 promotes association of Opa-negative, but not OpaD+, *N. gonorrhoeae* with human neutrophils in suspension. (A)** Primary human neutrophils in suspension were left untreated (UT) or treated with PMA. Neutrophils were exposed to anti-CD11b (44a; gray bar) or isotype control (white bar), then infected with CFSE-labeled Δopa Ngo. The percent of neutrophils with associated Ngo was calculated by imaging flow cytometry as described in **Fig. 2**. Data are the mean ± SEM for n=4 biological replicates. **(B-D)** Human neutrophils in suspension were left untreated (UT) or treated with PMA. Neutrophils were exposed to anti-CD11b (44a; gray bar) or isotype control (white bar), then infected with CFSE-labeled OpaD+ Ngo. The percent of neutrophils with associated **(B)** and internalized **(C)** bacteria was calculated as in **Fig. 1**. **(D)** has representative flow cytometry plots depicting CFSE (total) vs. DL650 (extracellular) fluorescence for OpaD+ Ngo; green = CFSE+ gate, blue = CFSE- gate, R1 = non-intact cells that are removed from analysis. Data are the mean ± SEM for 5 biological replicates, with statistical significance determined by two-way ANOVA followed by Tukey's multiple comparisons test. *** $P \leq 0.001$.

**Figure S9. Analysis of *M. smegmatis* phagocytosis by human neutrophils using imaging flow cytometry.** Adherent, IL-8 treated primary human neutrophils were infected with Tag-It Violet™ (TIV) labeled *M. smegmatis* (Msm) for 1 hr. Cells were fixed, stained with anti-human CD11b followed by anti-mouse IgG-AF488 secondary antibody, and analyzed by imaging flow cytometry. **(A-E)** Gating strategy: Single cells were identified by gating on cells with low area and high aspect ratio values **(A)**. Focused cells were identified by gating on objects in the single cell gate with high gradient RMS **(B)**. Neutrophils in the single, focused cell population were identified by CD11b positivity **(C)**. The population of neutrophils with associated Msm was quantified by gating on cells with ≥1 TIV spots (TIV+) **(D)**. The population of neutrophils with internalized Msm were identified by performing the TIV+ spot count for the area within a mask based on the CD11b+ surface staining (AF488 fluorescence area), eroded by 4 pixels **(E)**. **(F-G)** Representative images of neutrophils with intracellular Msm **(F)** and cell-associated but not internalized Msm **(G)**. The red mask in the bottom panel depicts the 4 pixel erosion mask based on surface CD11b fluorescence to distinguish intracellular from extracellular Msm.

#### Figure S1

**A**

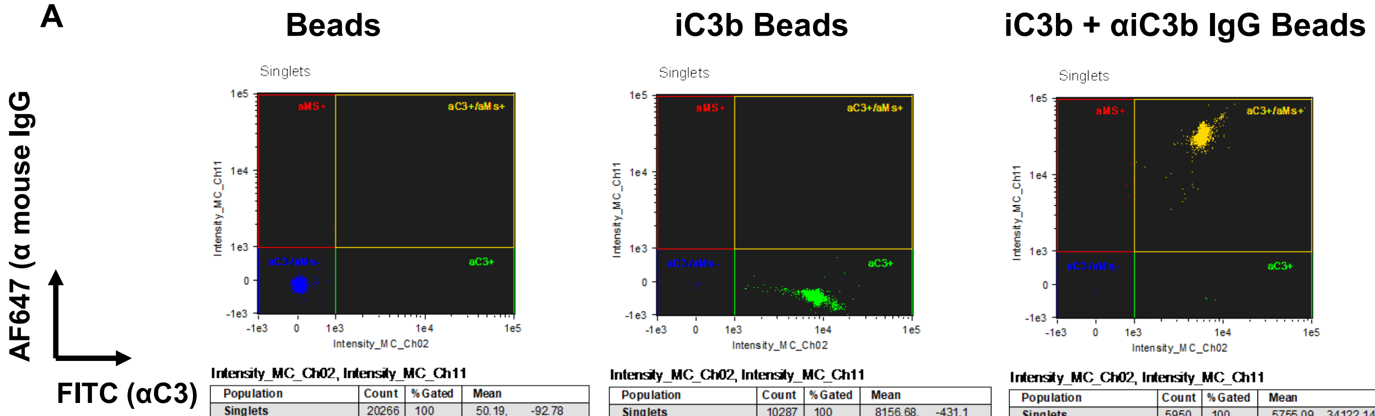


**Intensity_MC_Ch02, Intensity_MC_Ch11**

| Population | Count | %Gated | Mean |
|---|---|---|---|
| Singlets | 20266 | 100 | 50.19, -92.78 |
| aC3-/aMs- & Singlets | 20262 | 100 | 49.84, -102.34 |
| aMS+ & Singlets | 2 | 0.01 | 499, 1687.33 |
| aC3+ & Singlets | 1 | 0 | 6341.77, -304.67 |
| aC3+/aMs+ & Singlets | 0 | 0 | NaN, NaN |

**Intensity_MC_Ch02, Intensity_MC_Ch11**

| Population | Count | %Gated | Mean |
|---|---|---|---|
| Singlets | 10287 | 100 | 8156.68, -431.1 |
| aC3-/aMs- & Singlets | 7 | 0.07 | 42.78, -96.43 |
| aMS+ & Singlets | 0 | 0 | NaN, NaN |
| aC3+ & Singlets | 10273 | 99.9 | 8158.13, -431.95 |
| aC3+/aMs+ & Singlets | 1 | 0.01 | 43704.97, 8303.48 |

**Intensity_MC_Ch02, Intensity_MC_Ch11**

| Population | Count | %Gated | Mean |
|---|---|---|---|
| Singlets | 5950 | 100 | 5755.09, 34122.14 |
| aC3-/aMs- & Singlets | 5 | 0.08 | 359.9, 226.14 |
| aMS+ & Singlets | 6 | 0.1 | 834.44, 9308.63 |
| aC3+ & Singlets | 6 | 0.1 | 7406.38, -442.07 |
| aC3+/aMs+ & Singlets | 5933 | 99.7 | 5762.94, 34210.75 |

**B**

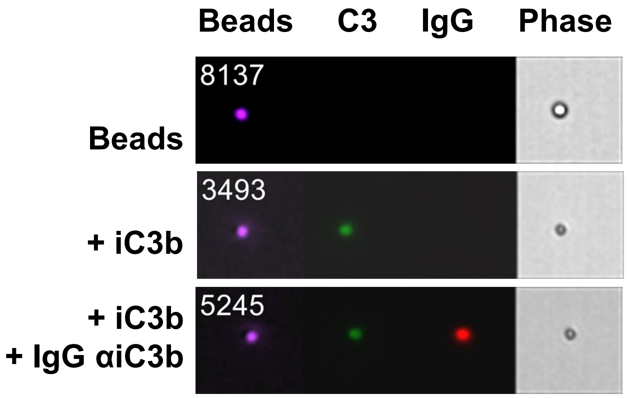

### Figure S2

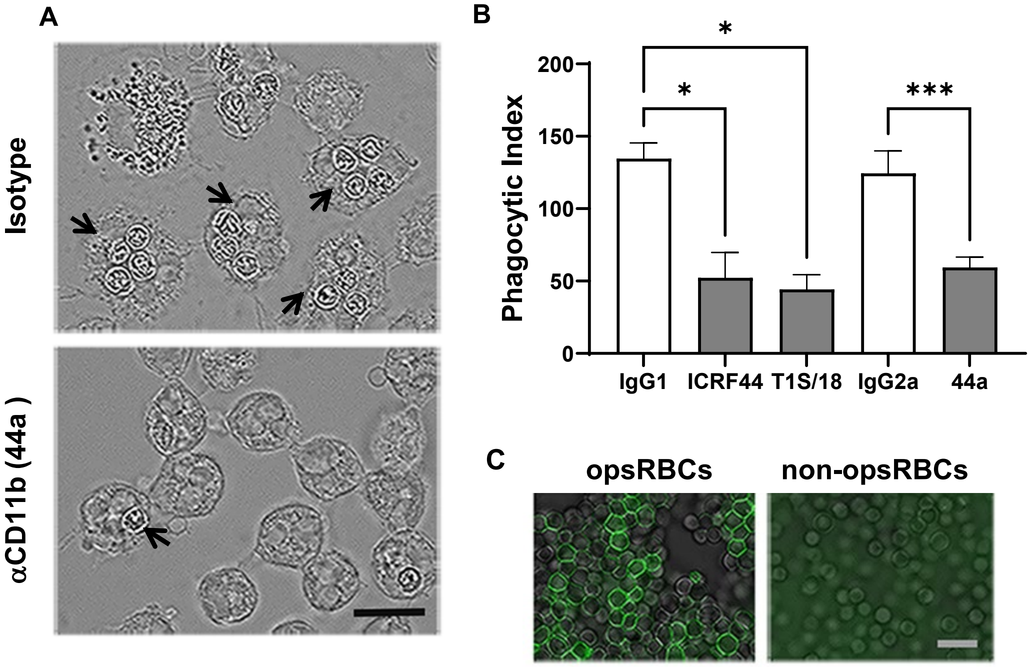
A
Isotype
αCD11b (44a )
B
Phagocytic Index
200
150
100
50
0
IgG1
ICRF44
T1S/18
IgG2a
44a
*
*
***
C
opsRBCs
non-opsRBCs

### Figure S3

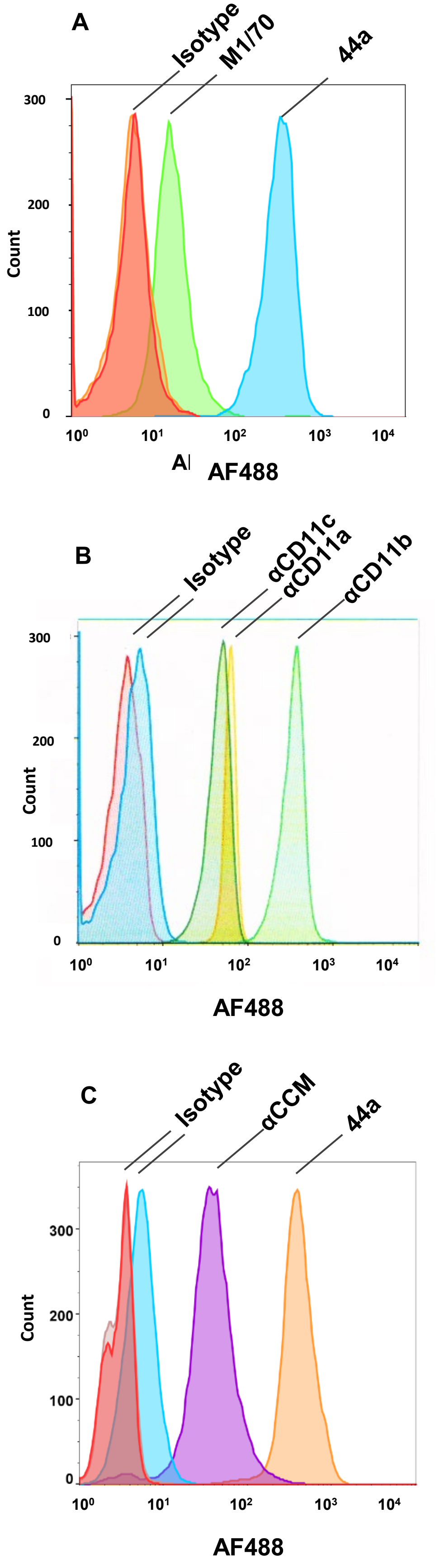
A
Isotype
M1/70
44a
300
200
100
0
Count
$10^0$
$10^1$
$10^2$
$10^3$
$10^4$
AF488
B
Isotype
αCD11c
αCD11a
αCD11b
300
200
100
0
Count
$10^0$
$10^1$
$10^2$
$10^3$
$10^4$
AF488
C
Isotype
αCCM
44a
300
200
100
0
Count
$10^0$
$10^1$
$10^2$
$10^3$
$10^4$
AF488

#### Figure S4

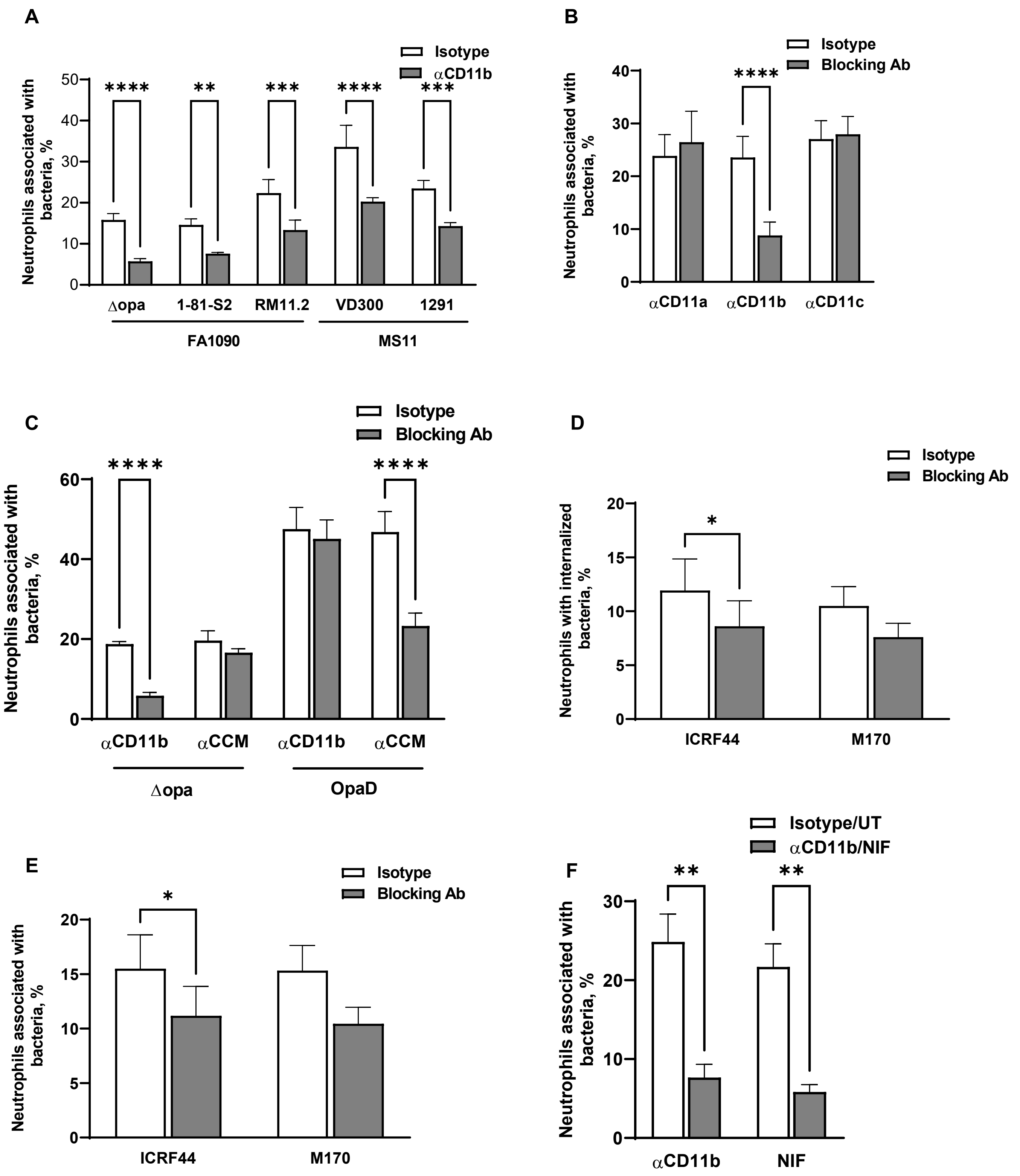
A
Isotype
αCD11b
Neutrophils associated with bacteria, %
0
10
20
30
40
50
****
**
***
****
***
Δopa
1-81-S2
RM11.2
VD300
1291
FA1090
MS11
B
Isotype
Blocking Ab
Neutrophils associated with bacteria, %
0
10
20
30
40
****
αCD11a
αCD11b
αCD11c
C
Isotype
Blocking Ab
Neutrophils associated with bacteria, %
0
20
40
60
****
****
αCD11b
αCCM
αCD11b
αCCM
Δopa
OpaD
D
Isotype
Blocking Ab
Neutrophils with internalized bacteria, %
0
5
10
15
20
*
ICRF44
M170
E
Isotype
Blocking Ab
Neutrophils associated with bacteria, %
0
5
10
15
20
*
ICRF44
M170
F
Isotype/UT
αCD11b/NIF
Neutrophils associated with bacteria, %
0
10
20
30
**
**
αCD11b
NIF

### Figure S5

## A

i

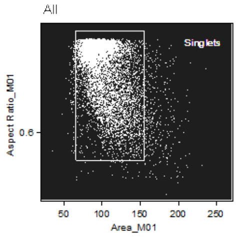


**Area_M01, Aspect Ratio_M01**

| Population | Count | %Gated | Mean |
|---|---|---|---|
| All | 14793 | 100 | 102.13, 0.8719 |
| Singlets | 14044 | 94.9 | 99.92, 0.881 |

ii

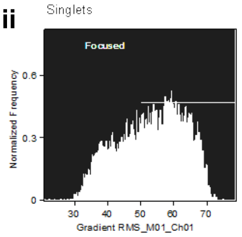


**Gradient RMS_M01_Ch01**

| Population | Count | %Gated | Mean |
|---|---|---|---|
| Singlets | 14044 | 100 | 53.15 |
| Focused & Singlets | 8881 | 63.2 | 59.46 |

iii

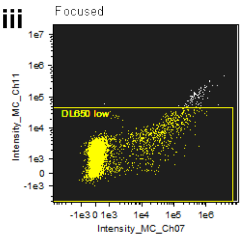


**Intensity_MC_Ch07, Intensity_MC_Ch11**

| Population | Count | %Gated | Mean |
|---|---|---|---|
| Focused & Singlets | 8881 | 100 | 16789.73, 4305.42 |
| DL650 low & Focused & Singlets | 8755 | 98.6 | 8112.21, 2566.24 |

iv

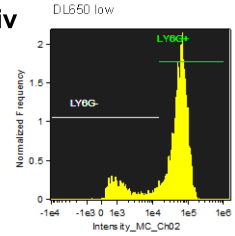


**Intensity_MC_Ch02**

| Population | Count | %Gated | Mean |
|---|---|---|---|
| DL650 low & Focused & Sing... | 8755 | 100 | 52699.97 |
| LY6G- & DL650 low & Focuse... | 1514 | 17.3 | 3384.21 |
| LY6G+ & DL650 low & Focuse... | 7240 | 82.7 | 62806.85 |

v

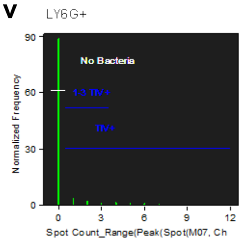


**Spot Count_Range(Peak(Spot(M07, Ch**

| Population | Count | %Gated | Mean |
|---|---|---|---|
| LY6G+ & DL650 low& Focuse... | 7240 | 100 | 0.3097 |
| No Bacteria & LY6G+ & DL65... | 6470 | 89.4 | 0 |
| TIV+ & LY6G+ & DL650 low &... | 770 | 10.6 | 2.912 |
| 1-3 TIV+ & LY6G+ & DL650 l... | 505 | 6.98 | 1.693 |

vi

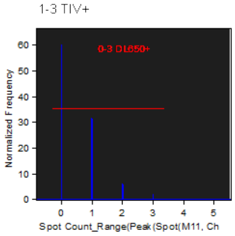


**Spot Count_Range(Peak(Spot(M11, Ch**

| Population | Count | %Gated | Mean |
|---|---|---|---|
| 1-3 TIV+ & LY6G+ & DL650 l... | 505 | 100 | 0.5188 |
| 0-3 DL650+ & 1-3 TIV+ & LY... | 503 | 99.6 | 0.501 |

vii

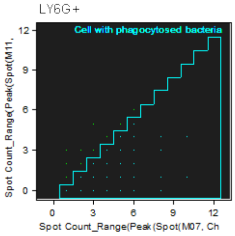


**Spot Count_Range(Peak(Spot(M07, Ch, Spot Count_Range(Peak(Spot(M11,**

| Population | Count | %Gated | Mean |
|---|---|---|---|
| LY6G+ & DL650 low & Focused... | 7240 | 100 | 0.3097, 0.09558 |
| Cell with phagocytosed bact... | 663 | 9.16 | 3.119, 0.7391 |

## B

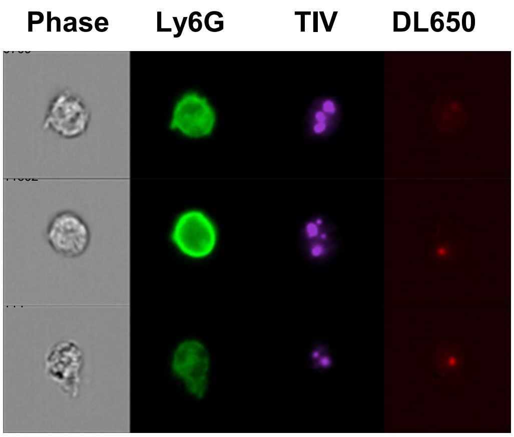


## C

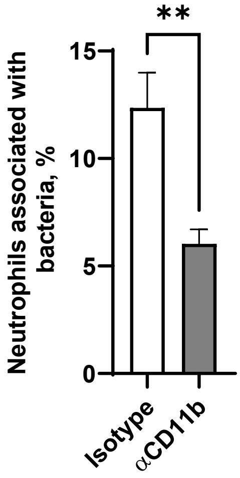


## D

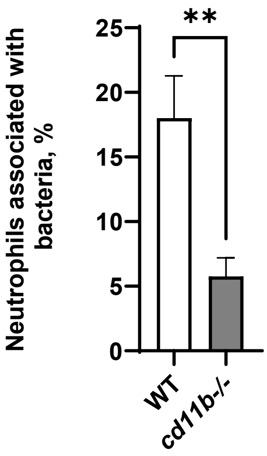

**Figure S6**

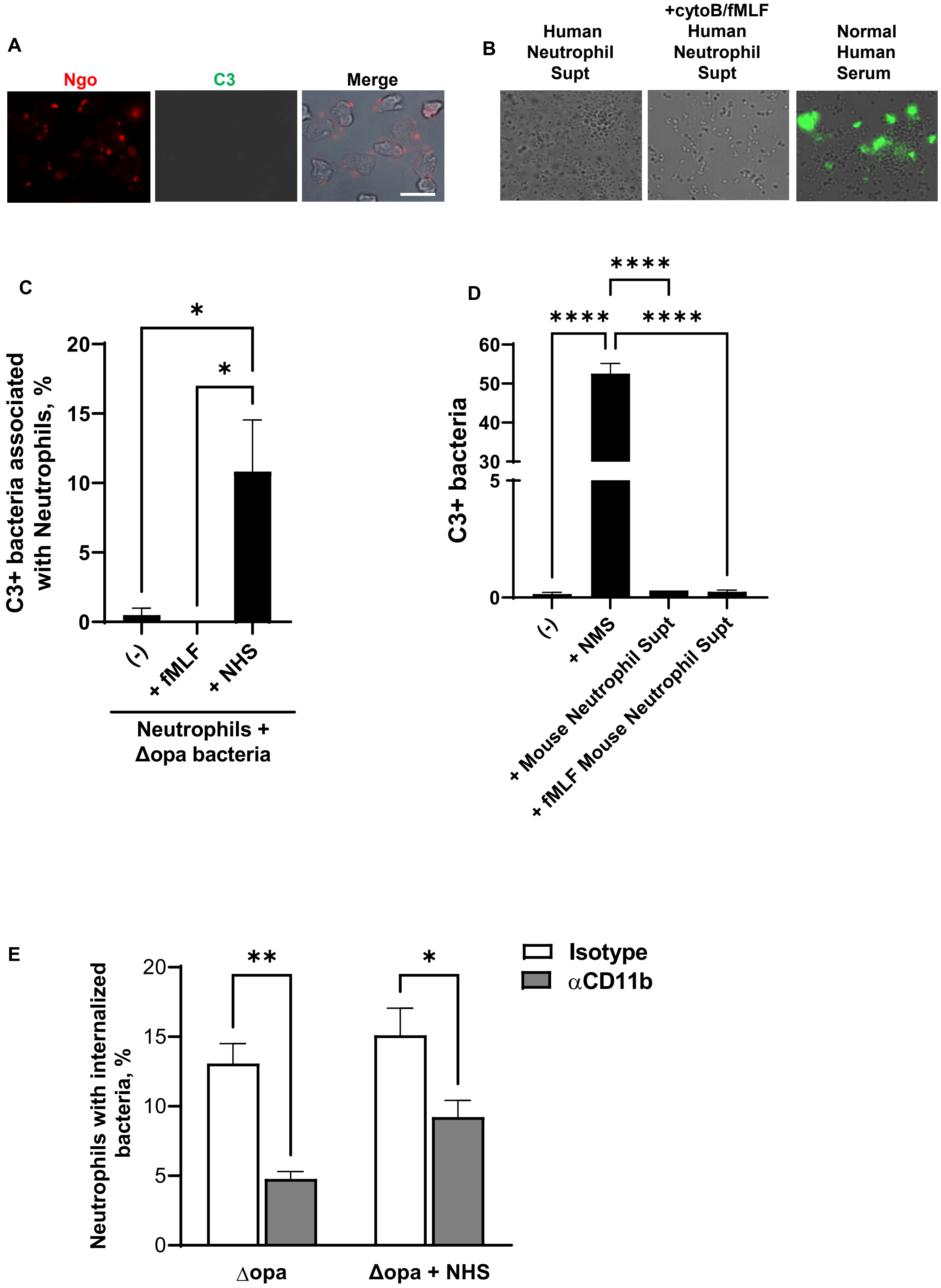
A
Ngo
C3
Merge
B
Human Neutrophil Supt
+cytoB/fMLF Human Neutrophil Supt
Normal Human Serum
C
C3+ bacteria associated with Neutrophils, %
20
15
10
5
0
*
*
(-)
+ fMLF
+ NHS
Neutrophils + Δopa bacteria
D
C3+ bacteria
60
50
40
30
5
0
****
****
****
(-)
+ NMS
+ Mouse Neutrophil Supt
+ fMLF Mouse Neutrophil Supt
E
Neutrophils with internalized bacteria, %
20
15
10
5
0
**
*
Isotype
αCD11b
Δopa
Δopa + NHS

### Figure S7

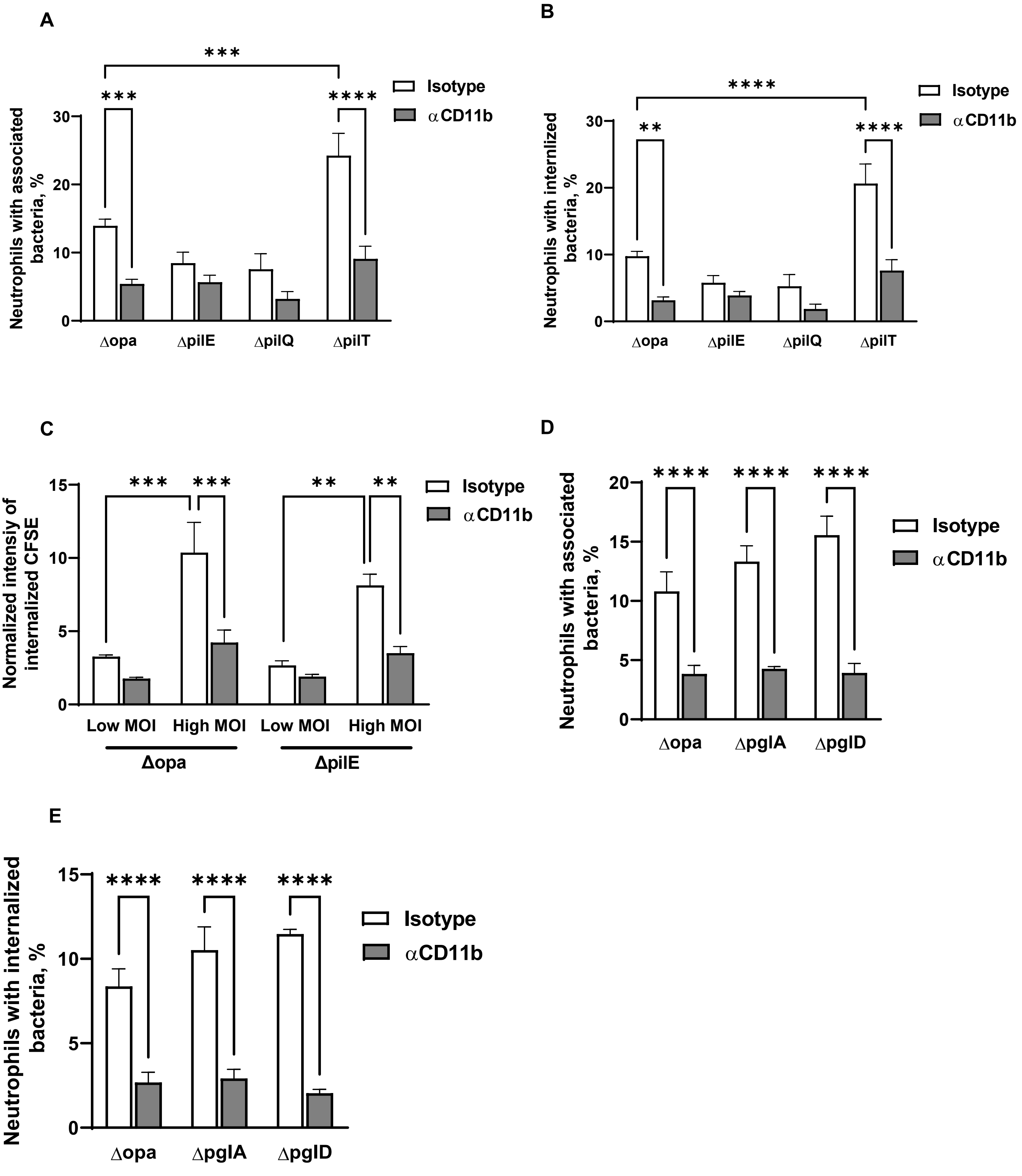
A
B
C
D
E
Neutrophils with associated bacteria, %
Neutrophils with internlized bacteria, %
Normalized intensiy of internalized CFSE
Neutrophils with internalized bacteria, %
Isotype
αCD11b
Δopa
ΔpilE
ΔpilQ
ΔpilT
Low MOI
High MOI
ΔpglA
ΔpglD
***
****
**

**Figure S8**

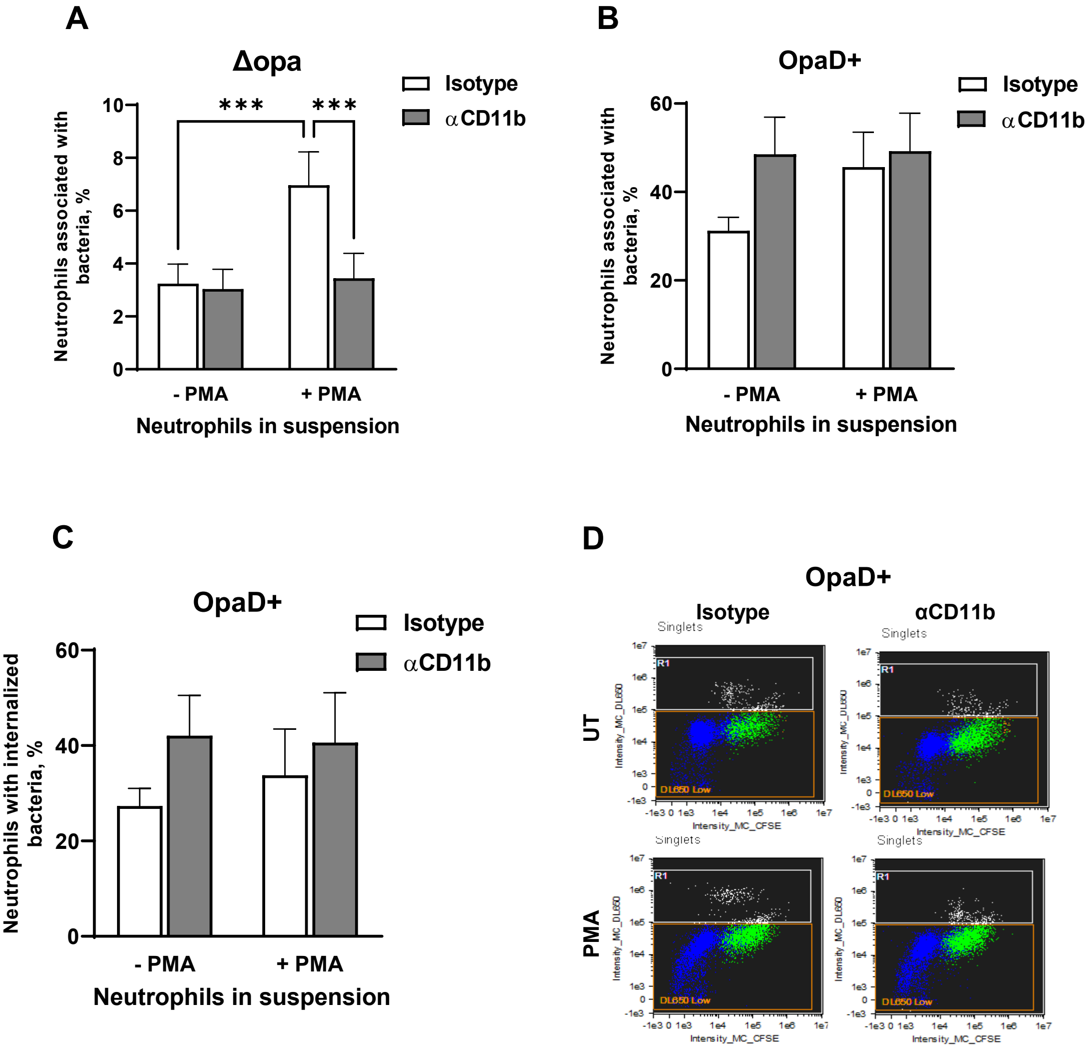
A
Δopa
Isotype
αCD11b
Neutrophils associated with bacteria, %
***
***
- PMA
+ PMA
Neutrophils in suspension
B
OpaD+
Isotype
αCD11b
Neutrophils associated with bacteria, %
- PMA
+ PMA
Neutrophils in suspension
C
OpaD+
Isotype
αCD11b
Neutrophils with internalized bacteria, %
- PMA
+ PMA
Neutrophils in suspension
D
OpaD+
Isotype
αCD11b
UT
PMA
Singlets
R1
DL650 Low
Intensity_MC_DL650
Intensity_MC_CFSE

### Figure S9

**A**

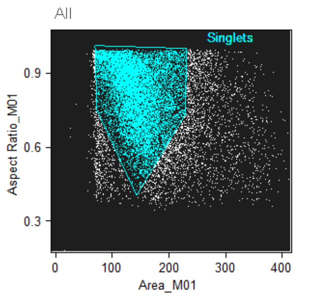


Area_M01, Aspect Ratio_M01

| Population | Count | % Gated | Mean |
|---|---|---|---|
| All | 13493 | 100 | 159.07, 0.7848 |
| Singlets | 10753 | 79.7 | 143.8, 0.8186 |

**B**

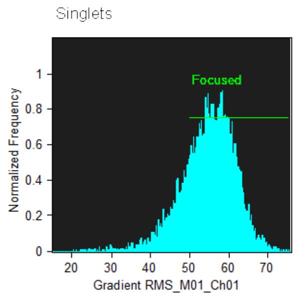


Gradient RMS_M01_Ch01

| Population | Count | % Gated | Mean |
|---|---|---|---|
| Singlets | 10753 | 100 | 54.81 |
| Focused & Singlets | 8487 | 78.9 | 57.38 |

**C**

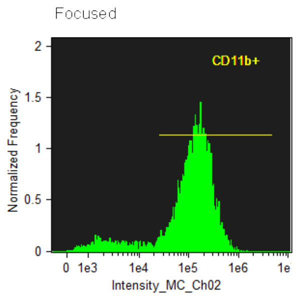


Intensity_MC_Ch02

| Population | Count | % Gated | Mean |
|---|---|---|---|
| Focused & Singlets | 8487 | 100 | 154485.02 |
| CD11b+ & Focused & Singlets | 7502 | 88.4 | 173865.06 |

**D**

**Neutrophils with associated Msm**

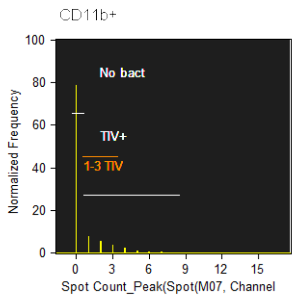


Spot Count_Peak(Spot(M07, Channel

| Population | Count | % Gated | Mean |
|---|---|---|---|
| CD11b+ & Focused & Singlets | 7502 | 100 | 0.5032 |
| No bact & CD11b+ & Focused... | 5939 | 79.2 | 0 |
| TIV+ & CD11b+ & Focused & ... | 1559 | 20.8 | 2.397 |
| 1-3 TIV & CD11b+ & Focused... | 1236 | 16.5 | 1.744 |

**E**

**Neutrophils with internalized Msm**

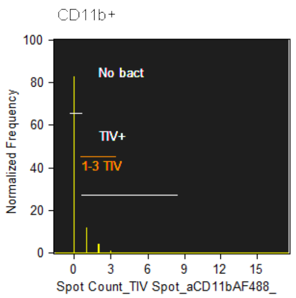


Spot Count_TIV Spot_aCD11bAF488_

| Population | Count | % Gated | Mean |
|---|---|---|---|
| CD11b+ & Focused & Singlets | 7502 | 100 | 0.2333 |
| No Bact Internal | 6237 | 83.1 | 0 |
| TIV+ Internal | 1265 | 16.9 | 1.383 |
| 1-3 TIV Internal | 1248 | 16.6 | 1.348 |

**F**

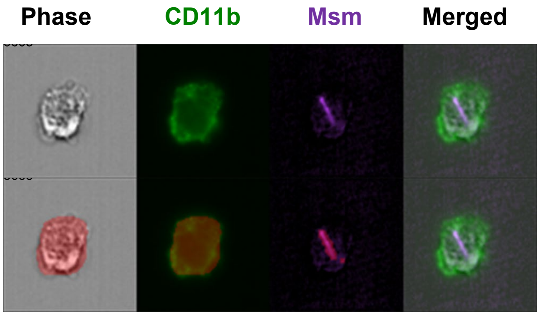


**G**

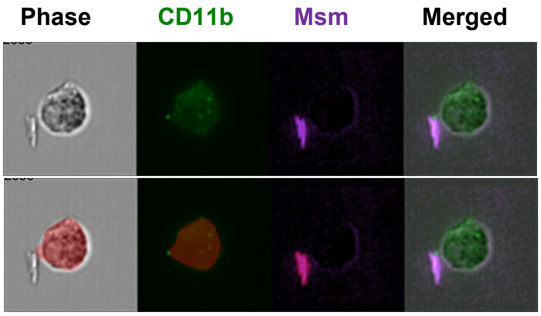
